# Nuclear MTHFD2 secures mitosis progression by preserving centromere integrity

**DOI:** 10.1101/2023.06.01.543193

**Authors:** Natalia Pardo-Lorente, Anestis Gkanogiannis, Luca Cozzuto, Antoni Gañez Zapater, Lorena Espinar, Laura García-López, Rabia Gül Aydin, Evangelia Darai, Jacqueline Severino, Laura Batlle-Morera, Julia Ponomarenko, Sara Sdelci

## Abstract

Subcellular compartmentalization of metabolic enzymes may elicit specific cellular functions by establishing a unique metabolic environment. Indeed, the nuclear translocation of certain metabolic enzymes is required for epigenetic regulation and gene expression control. Here, we reveal that, in cancer cells, the mitochondrial enzyme methylenetetrahydrofolate dehydrogenase 2 (MTHFD2) localizes in the nucleus during the G2-M phase of the cell cycle to secure mitosis progression. Nuclear MTHFD2 interacts with proteins involved in mitosis regulation and centromere stability, including the methyltransferases KMT5A and DNMT3B. Loss of MTHFD2 induces centromere overexpression and severe methylation defects and impedes correct mitosis completion. As a consequence, MTHFD2 deficient cells accumulate chromosomal aberrations arising from chromosome congression and segregation defects. Blocking the catalytic nuclear function of MTHFD2 recapitulates the phenotype observed in MTHFD2 deficient cells, attributing to nuclear MTHFD2 an enzymatic active role in controlling mitosis. Our discovery uncovers a nuclear moonlighting role for the cancer target MTHFD2, and emphasizes that cancer metabolism rewiring may encompass the relocation of metabolic enzymes to alternative subcellular compartments.

## Introduction

Enzymes of central metabolism can translocate to the cellular nucleus, impacting chromatin remodeling, epigenetics and transcription regulation^1–5^. In cancer, clear examples of how nuclear metabolism can interact with epigenetics are given by the control of histone acetylation and methylation *via* the nuclear translocation of enzymes responsible for the synthesis of the required substrates and cofactors^6–9^.

One-carbon folate metabolism is a pivotal pathway in cancer progression, being indispensable for the *de novo* synthesis of nucleotides, amino acid homeostasis, DNA and histone methylation, and the maintenance of the cellular redox state^10^. Folate metabolism is a compartmentalized pathway, primarily localized in the cytosol and the mitochondria. In contrast to their cytoplasmic counterparts, in cancer, mitochondrial one-carbon metabolism enzymes are greatly upregulated, making them central to tumor growth^11, 12^.

The mitochondrial one-carbon metabolism enzyme methylenetetrahydrofolate dehydrogenase 2 (MTHFD2) emerged as the most consistently metabolic gene overexpressed in tumors^13^. High levels of MTHFD2 are associated with a worse outcome in several cancer types, including breast^14^, colon^15^ and lung^16^ cancer. MTHFD2 has been shown to support cell proliferation and survival *in vitro* and tumor growth *in vivo*^15–19^, and to promote metastatic features such as cell migration and invasion^17, 20^. The fact that MTHFD2 is lowly expressed in adult tissue but overexpressed in cancer^13^, made this enzyme very attractive for the development of novel, metabolism-centered cancer therapies^21–23^. Despite its canonical mitochondrial role, MTHFD2 can also localize in the cellular nucleus^24, 25^, and we previously showed that it is bound to chromatin^26^. However, the nuclear function of MTHFD2 has been poorly characterized and remains elusive.

Cancer cells are immortal and hence indefinitely continue to duplicate their genetic material to distribute it into their daughter cells through the accurate process of mitosis. Even though cancer cells accumulate mutations that provide fitness advantages, mitotic defects that lead to macroscopic genetic aberrations are deleterious and often cause cell death^27^. For this reason, drugs that affect mitosis progression, such as microtubule-targeting agents, are commonly used as anticancer therapy^28^. Several mechanisms ensure faithful chromosome segregation, and centromeres are important pillars in this process. Centromeres orchestrate the chromosomal attachment to spindle fibers through the assembly of the kinetochore complex and sustain sister chromatid cohesion by assisting in the Cohesin complex formation^29–31^. Centromeric and peri-centromeric DNA is compacted into constitutive heterochromatin and is enriched in Cohesin, Condensin and Topoisomerase II^32^. Heterochromatin marks H3K9me3, H3K27me3 and H4K20me1 decorate the centromere, contributing to chromatin compaction and the formation of the kinetochore^33–36^. To ensure a correct cell division, equally important is to maintain DNA methylation. Perturbation of DNA methylation has been associated with mitotic defects and genomic instability^37–40^. DNA methylation at centromeres is highly abundant, and it has been proposed to maintain chromatin structure and prevent chromosome segregation errors and genomic instability^41^. Finally, controlling the expression of the centromeric region is key to ensuring the proper assembly of the kinetochore complex. The expression of centromeric non-coding alpha-satellite RNAs regulates the loading of the centromeric histone variant CENP-A and contributes to the recruitment of inner kinetochore proteins to the centromeric region^42^.

Here, for the first time, we reveal that, in cancer cells, the enzyme MTHFD2 localizes in the nucleus to regulate DNA and centromeric histone methylation, and consequently ensure accurate mitotic division. After validating that MTHFD2 is chromatin-bound in a variety of cancer cell lines, we investigated its nuclear interactome and discovered that MTHFD2 nuclear partners are mostly cell cycle regulators involved in centromere and kinetochore stability and mitosis progression. Interestingly, among MTHFD2 nuclear interactors we retrieved methyltransferases responsible for depositing methylation marks at centromeres, such as KMT5A^33, 43^, DNMT3B^44^ and PRMT1^45^. The absence of MTHFD2 leads to a drastic reduction of DNA and centromeric histone methylation, increased centromeric alpha-satellite expression and accumulation of genomic aberrations. Cell cycle progression is also impaired when cells lack MTHFD2, with a significant reduction of mitotic events. When analyzing specific mitotic phases, we discovered that the absence of MTHFD2 induces chromosome congression and segregation defects, which result in the accumulation of micronuclei. Importantly, the inhibition of the nuclear function of MTHFD2 recapitulates this phenotype, attributing to the nuclear localization of MTHFD2 an enzymatic active role controlling centromeric heterochromatin maintenance and correct mitotic cell division.

## Results

### MTHFD2 is recruited to chromatin across cancer types

A meta-analysis comprising microarray expression data covering 19 types of tumors highlighted the mitochondrial folate enzyme MTHFD2 as the top-scoring upregulated metabolic enzyme in cancer^13^. To corroborate this finding, we retrieved RNA-sequencing data from The Cancer Genome Atlas (TCGA) database^46^. We kept those solid tumor types where paired normal tissue data were available (Extended Data Fig. 1a), and we confirmed that *MTHFD2* was significantly upregulated in 13 from 15 tumor types, being breast carcinoma, colon adenocarcinoma, lung adenocarcinoma and lung squamous cell carcinoma the most significant classes (Fig. 1a). Then, we asked whether MTHFD2 expression alone could be used to predict the status of a sample (tumor versus healthy) in breast, lung and colon cancer. By training a random forest machine learning algorithm with a subset of the expression data, we obtained values of accuracy of prediction using the unseen data higher than 0.84, and values of area under the curve (AUC) of the receiver operating curve (ROC) between 0.77 and 0.88 (Extended Data Fig. 1b-d), confirming the predictive usefulness of MTHFD2 expression.

**Fig. 1:**
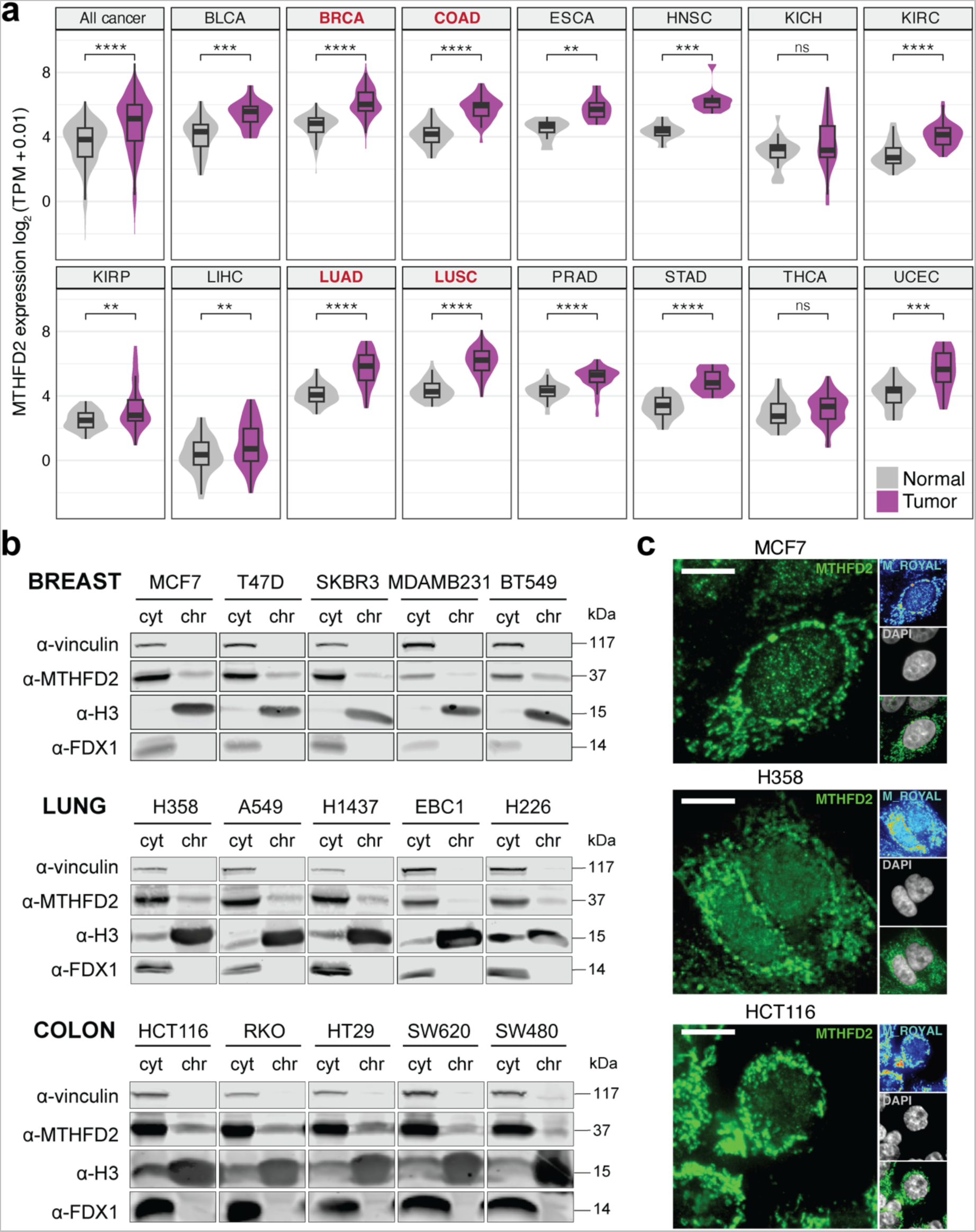
MTHFD2 localizes on chromatin in cancer cells. **a**, Comparison between Transcripts per Million (TPM) expression values of MTHFD2 in healthy and tumor tissues; paired two-tailed Wilcoxon test (ns, non-significant; **, *P*<0.01; ***, *P*<0.001; ****, *P*<0.0001). BLDA, bladder urothelial carcinoma; BRCA, breast invasive carcinoma; COAD, colon adenocarcinoma; ESCA, esophageal carcinoma; HNSC, head and neck squamous cell carcinoma; KICH, kidney chromophobe; KIRC, kidney renal clear cell carcinoma; KIRP, kidney renal papillary cell carcinoma; LIHC, liver hepatocellular carcinoma; LUAD, lung adenocarcinoma; LUSC, lung squamous cell carcinoma; PRAD, prostate adenocarcinoma; STAD, stomach adenocarcinoma; THCA, thyroid carcinoma; UCEC, uterine corpus endometrial carcinoma. **b,** Western blot of cytosolic (cyt) and chromatin (chr) fractions after subcellular fractionation in a panel of cancer cell lines. Vinculin, histone H3 and FDX1 are used as cytosolic, nuclear and mitochondrial markers, respectively. **c,** Immunofluorescence of MTHFD2 in MCF7 (up), H358 (middle) and HCT116 (down) cells. MTHFD2 is shown in green (left) or royal (right) and DAPI in grey; confocal mode, scale bar 10 μm.

Although MTHFD2 is primarily found in the mitochondria, its nuclear localization was first reported in 2015^25^. A few years later, we reported that MTHFD2, along with other members of the one-carbon metabolism, localizes on chromatin^26^. Since our gene expression meta-analysis supported that MTHFD2 is highly relevant in breast, colon and lung cancer, we chose a panel of cancer cell lines belonging to these cancer types to address whether MTHFD2 was recurrently recruited to chromatin. This panel comprised 5 breast cancer cell lines of different subtypes (hormone receptor-positive: MCF7 and T47D; HER2-positive: SKBR3; and triple-negative: MDAMB231 and BT549), 5 colon cancer cell lines of varied subtypes (consensus molecular subtype (CMS) 1: RKO; CMS2: SW480 and SW620; CMS4: HCT116; and HT29) and 5 lung cancer cell lines belonging to both lung adenocarcinoma (A459, H358, H1437) and lung squamous cell carcinoma (EBC1, H226). Retrieving data from the Cancer Cell Line Encyclopedia (CCLE)^47^, we observed that MTHFD2 RNA expression level was generally high across all these cell lines, while protein expression level was more variable (Extended Data Fig. 1e,f). Moreover, MTHFD2 was not essential in any of the cell lines (Extended Data Fig. 1g). With these cell lines, we performed a subcellular fractionation to isolate their proteome associated with the chromatin (i.e., chromatome). After verifying the purity of the fractions by blotting Vinculin, H3 and FDX1 as cytosolic, nuclear and mitochondrial markers, respectively, MTHFD2 could be detected on chromatin in all the cancer cell lines tested (Fig. 1b). Confocal immunofluorescence with the cell lines showing the highest MTHFD2 chromatin abundance in Western blot from each cancer type confirmed MTHFD2 nuclear localization (Fig. 1c). Additionally, we performed confocal microscopy on colon cancer patient-derived organoids, where we also observed a mitochondrial and nuclear MTHFD2 signal, again corroborating MTHFD2 nuclear localization (Extended Data Fig. 1h).

Our data indicate that the folate enzyme MTHFD2 is not only recurrently upregulated in cancer but also that its expression level can predict whether a tissue is cancerous. Besides, we showed that MTHFD2 consistently localizes on chromatin across cancer cell types and patient-derived organoids, which may imply a relevant contribution of nuclear MTHFD2 in cancer progression.

### Nuclear MTHFD2 interacts with mitotic proteins

To unravel which could be the functionality of nuclear MTHFD2 in cancer, we characterized the nuclear MTHFD2 interactome using the HCT116 cell line as a model. We selected this cell line because the levels of MTHFD2 in chromatin are high (Fig. 1b), it is a diploid cell line and it has been previously used to study the role of MTHFD2 in cancer^15^. Besides, among the colon cancer cell lines, it has the highest RNA and protein MTHFD2 expression and the highest MTHFD2 dependency (Extended Data Fig. 1e-g).

With this purpose, we coupled a pull-down of MTHFD2 in the cytosolic and chromatin fractions, obtained after subcellular fractionation, to mass spectrometry (MS) (Supplementary Table 1). We verified the purity of the fractions by checking the relative abundance of the detected proteins in the different subcellular compartments using Human Protein Atlas (HPA)^48^ annotation. As expected, cytosolic MTHFD2 and IgG immunoprecipitations (IP) spanned a wider range of cell compartments, with the cytosol with the highest proportion (41 and 43%, respectively). On the other hand, chromatin MTHFD2 and IgG IPs contained mainly nuclear proteins (80 and 85%, respectively) (Extended Data Fig. 2a). Then, we prioritized a list of interactors using the Significance Analysis of INTeractome (SAINT) software^49^. We identified 127 and 43 potential MTHFD2 interactors in the cytosolic and chromatin fractions, respectively (Extended Data Fig. 2b, Supplementary Table 2). Using the HPA annotation for subcellular localization, we categorized MTHFD2 interactors according to their nuclear localization. From the cytosolic interactors, 35% of the proteins were classified as not nuclear, 36% as mainly nuclear and 12% as additionally nuclear, while the localization of the rest was not available. From the chromatin interactors, 21% were not nuclear, 51% were mainly nuclear and the rest were not available (Extended Data Fig. 2c). Thus, although the proportion of nuclear proteins was higher in the chromatin fraction, it was also quite noticeable in the cytosol. We therefore considered as nuclear MTHFD2 interactors those which fell in the categories of mainly nuclear or additionally nuclear, from both fractions (Supplementary Table 2).

**Fig. 2:**
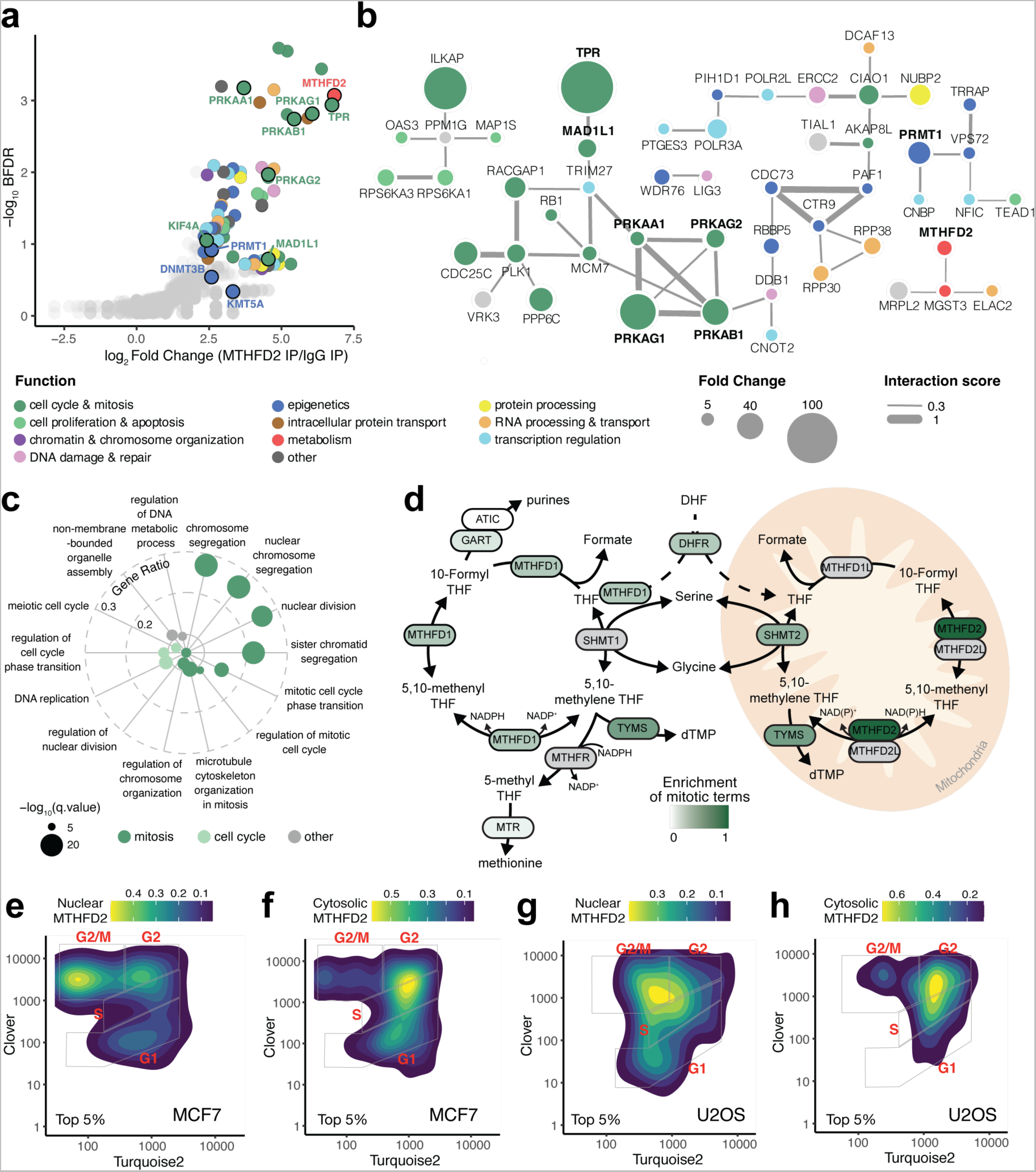
Nuclear MTHFD2 interacts with key mitotic proteins. **a**, Volcano plot of top nuclear MTHFD2 interactors identified in HCT116 cells. Interactors with a log_2_ fold change >= 2.3 and Bonferroni False Discovery Rate (BFDR) <= 0.2 are colored according to their functional category. **b,** Network of top nuclear MTHFD2 interactors. The color indicates the functional category, the size represents the fold change and the width of the edges shows the interaction score. **c,** Biological Process gene ontologies enriched in MTHFD2 core co-expressed genes. **d,** One-carbon folate metabolism pathway with enzymes colored by the enrichment of mitotic terms in the Gene Ontology enrichment analysis performed with their core co-expressed genes. Gray indicates the absence of data. DHF, dihydrofolate; THF, tetrahydrofolate. Scheme adapted from Lin *et al*^4^. **e-h,** Two-dimensional density plots of 5% top cells with the highest nuclear (e,g) or cytosolic (f,h) MTHFD2 signal obtained from FUCCI4 adapted MCF7 (e,f) and U2OS (g,h) cells along with the cell cycle phase; turquoise and clover mean intensities (x and y axis, respectively) are in log_10_ scale.

To get insights into the role of nuclear MTHFD2, we functionally categorized its top nuclear interactors (Fig. 2a, Extended Data Fig. 2d). The most abundant functional category in our interactome was cell cycle and mitosis, with 23% of the top interactors, followed by epigenetics and transcription regulation (Extended Data Fig. 2e). IntAct^50^ network analysis showed a high degree of connectivity, with previously described interactions among 57% of our hits (Fig. 2b). The only known nuclear interactor of MTHFD2 was Microsomal Glutathione S-Transferase 3 (MGST3) and hence, our data provides an entirely novel set of nuclear MTHFD2 partners. The top MTHFD2 nuclear interactor was Nucleoprotein TPR, a nucleopore complex protein that stabilizes the mitotic spindle assembly checkpoint proteins MAD1L1 and MAD2L1 during mitosis, contributing to the activation of the spindle assembly checkpoint^51^. MAD1L1 was also among the MTHFD2 interactors (Fig. 2a,b). Additionally, we retrieved all the members of the AMP-activated protein kinase (AMPK) complex (PRKAA1, PRKAB1, PRKAG1 and PRKAG2), which is known to be involved in mitosis progression^52^, anaphase spindle length determination^53^ and spindle orientation^54^. Kinesin KIF4A, a substrate of AMPK^53^ that participates in chromosome condensation and segregation^55^, and Protein Arginine N-Methyltransferase 1 (PRMT1), which methylates Inner Centromere Protein (INCENP) triggering the centromeric recruitment and activation of AURKB^45^, also scored as significant MTHFD2 nuclear interactors. Furthermore, we identified N-Lysine Methyltransferase KMT5A, which deposits H4K20me1 at centromeric CENP-A containing nucleosomes enabling kinetochore assembly^33, 43, 56^, and DNA Methyltransferase 3B (DNMT3B), which is recruited to the centromeric and pericentromeric regions by CENP-C for DNA methylation^44^, as putative MTHFD2 nuclear interactors. MTHFD2 pull-down experiments performed in HTC116 and H358 nuclear extracts (Extended Data Fig. 2f) validated most of these interactions (Extended Data Fig. 2g,h).

Since co-expression can be used to infer functionality, we exploited the TCGA RNA-sequencing data to identify genes that are co-expressed with MTHFD2. We selected as positively correlated genes those that were co-expressed with MTHFD2 in at least 10 different cancer types, and we performed an over-representation analysis of Gene Ontology (GO) Biological Processes (Extended Data Fig. 3a,b). Top-scoring terms were mostly related to mitosis and cell cycle (Fig. 2c), showing a particular enrichment for terms associated with chromosome segregation. We repeated such analysis with all the enzymes of the folate pathway (Supplementary Table 3,4) and we obtained the proportion of significant terms related to mitosis or the cell cycle (Extended Data Fig. 3c,d). MTHFD2 was by far the folate enzyme with the highest proportion of mitotic-related (Fig. 2d) and cell cycle-related terms (Extended Data Fig. 3e), followed by TYMS. A similar analysis was done at the protein level with ProteomeHD^57^, which uses proteomics data in response to biological perturbations to perform co-regulation analysis using unsupervised machine learning. We identified the 5% strongest co-regulated proteins with MTHFD2, and an over-representation analysis of GO Biological Processes using such co-regulated proteins revealed again several cell cycle and mitotic terms (Extended Data Fig. 3f).

**Fig. 3:**
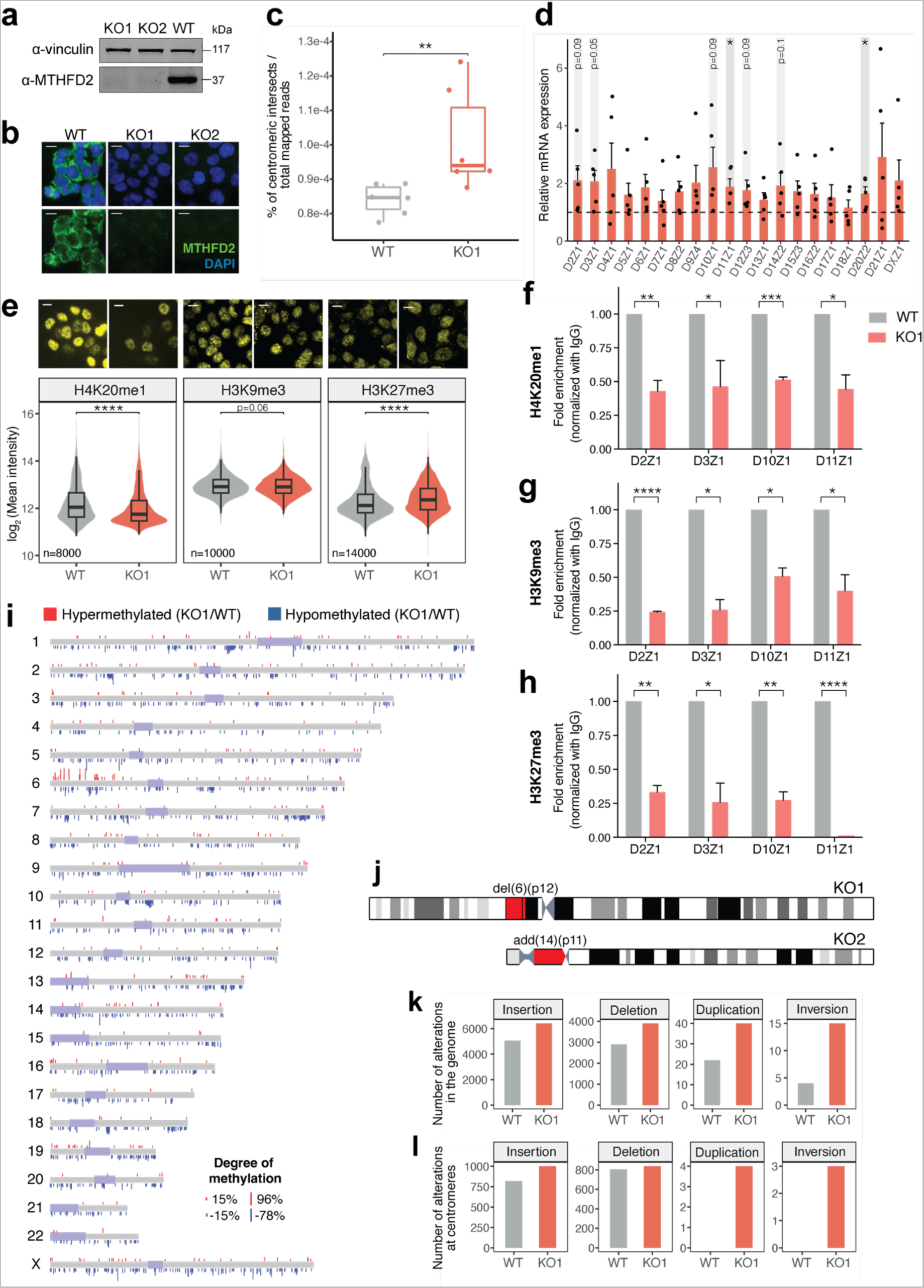
MTHFD2 KO induces aberrant centromere overexpression, strong methylation defects and increased structural variation. **a**, Western blot of HCT116 MTHFD2 knock-out (KO1, KO2) and wild-type (WT) cells. Vinculin is used as loading control. **b,** Immunofluorescence of MTHFD2 WT, KO1 and KO2 cells. MTHFD2 is shown in green and DAPI in blue; non-confocal mode, scale bar 10 μm. **c,** Percentage of centromeric intersects normalized by the total mapped reads in WT and KO1 cells; unpaired two-tailed Wilcoxon test (**, *P*<0.01). **d,** Relative mRNA expression of 20 chromosomal centromeres in KO1 cells normalized to WT cells. The dashed line indicates fold change = 1; means + s.e. (*n*=5), one-sample two-tailed *t-*test (*, *P*<0.05). **e,** Comparison of the log_2_ mean intensity of nuclear levels of histone marks H4K20me1, H3K9me3 and H3K27me3 in WT and KO1 cells; unpaired two-tailed Wilcoxon test (ns, non-significant; ****, *P*<0.0001). Representative images are shown above; non-confocal mode, scale bar 10 μm. **f-h,** Fold enrichment of H4K20me1 (f), H3K9me3 (g) and H3K27me3 (h) signal normalized to IgG in centromeric regions of 4 independent chromosomes in WT and KO1 cells; means + s.d. (*n*=3), one-sample two-tailed *t-*test (*, *P*<0.05; **, *P*<0.01; ***, *P*<0.001; ****, *P*<0.0001). **i,** Whole-genome scheme showing the hypermethylated CpG sites in red and hypomethylated CpG sites in blue. Regions shown in pale purple correspond to the centromeres and peri-centromeres. The height of the bars is proportional to the degree of hyper- or hypomethylation. **j,** Chromosomes 6 and 14 showing the peri-centromeric alterations found in KO1 and KO2 cells in red. **k-l,** Number of alterations (insertions, deletions, duplications or inversions) found in WT and KO1 cells in the whole-genome (k) or at the centromeric regions (l).

Given the possible implication of MTHFD2 in mitosis, we investigated whether its nuclear localization is cell cycle-dependent. We first explored whether global MTHFD2 protein levels differed in cell cycle stages using the data published by Ly and colleagues^58^, which indicated a significantly higher amount of MTHFD2 in the G2 phase of the cell cycle (Extended Data Fig. 4a). We then employed a FUCCI4^59^ adapted MCF7 (Extended Data Fig. 4b) and U2OS (Extended Data Fig. 4c) cell line reporters that allow cell cycle phase tracking, and we observed that cells in the G2-M phase of the cell cycle had the highest levels of nuclear MTHFD2 (Fig. 2e,g), which strengthens the hypothesis of an MTHFD2 nuclear role controlling mitosis. Cells with the highest cytosolic MTHFD2 levels were instead in the S-G2 phase of the cell cycle (Fig. 2f,h).

**Fig. 4:**
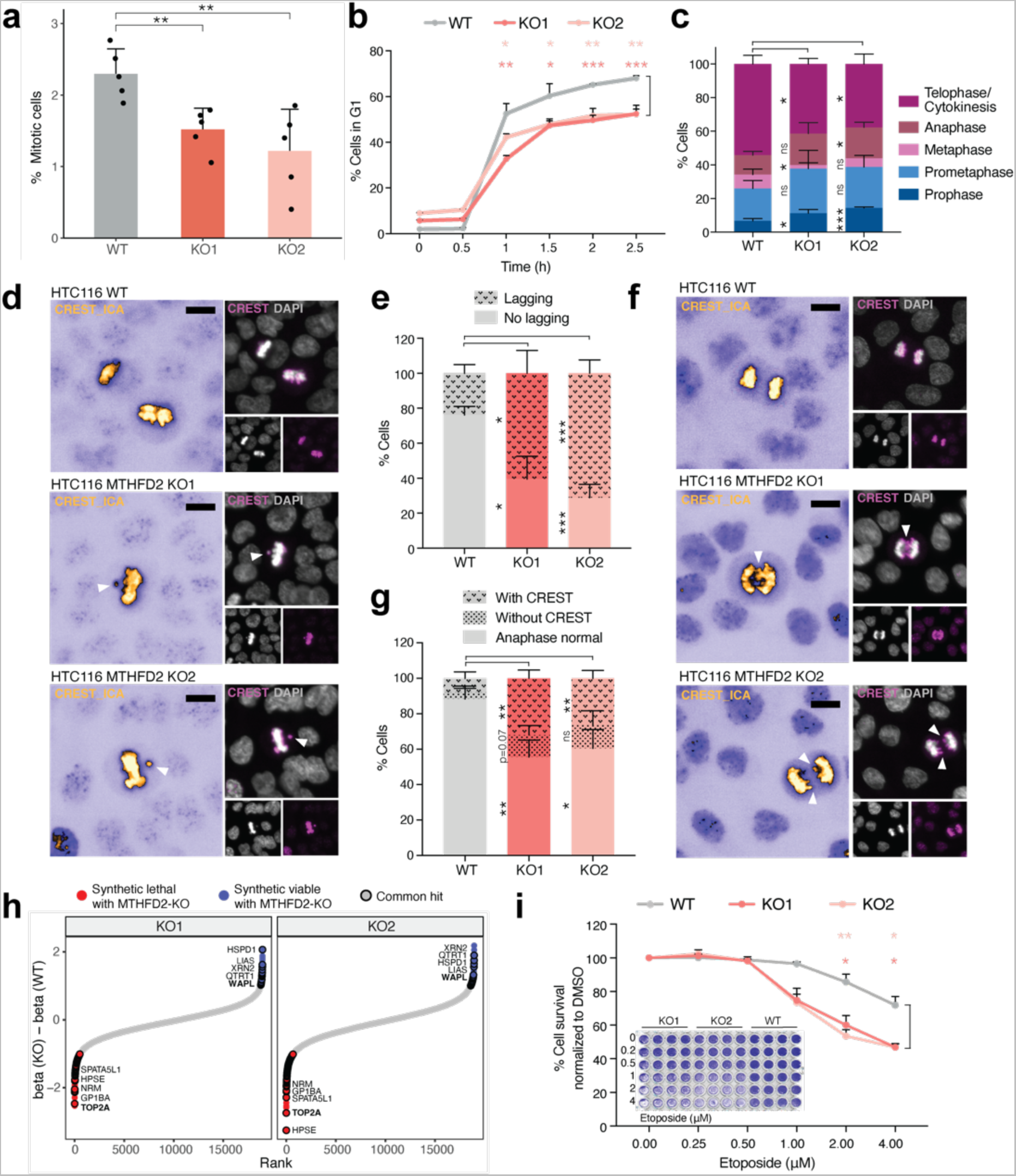
MTHFD2 loss impairs mitosis progression. **a**, Percentage of mitotic cells in HCT116 MTHFD2 wild-type (WT) and knock-out (KO1, KO2) conditions, measured with the mitotic marker histone H3 phospho-Ser10 by high-throughput immunofluorescence; means + s.d. (*n*=5), unpaired two-tailed *t*-test (**, *P*<0.01). **b,** Percentage of WT, KO1 and KO2 cells in G1 phase at 0, 0.5, 1, 1.5, 2 and 2.5-hour release after RO-3306 drug treatment for 20 hours; means + s.d. (*n*=3), at indicated times, unpaired two-tailed *t*-test (*, *P*<0.05; **, *P*<0.01; ***, *P*<0.001). **c,** Percentage of WT, KO1 and KO2 cells at different mitotic phases; means + s.d. (*n*=3), a minimum of 150 mitotic cells per replicate were analyzed, unpaired two-tailed *t*-test (ns, non-significant; *, *P*<0.05; ***, *P*<0.001). **d-e,** Representative images (d) and quantification (e) of lagging chromosomes in metaphase in WT, KO1 and KO2 cells. CREST is shown in ICA (left) or magenta (right) and DAPI in grey; non-confocal mode, scale bar 10 μm. For the quantification, means + s.d. (*n*=3), a minimum of 10 metaphase cells were analyzed, unpaired two-tailed *t*-test (*, *P*<0.05; ***, *P*<0.001). **f-g,** Representative images (f) and quantification (g) of anaphase defects in WT, KO1 and KO2 cells. CREST is shown in ICA (left) or magenta (right) and DAPI in grey; non-confocal mode, scale bar 10 μm. For the quantification, means + s.d. (*n*=3), a minimum of 25 anaphase cells were analyzed, unpaired two-tailed *t*-test (ns, non-significant; *, *P*<0.05; **, *P*<0.01). **h,** Difference between the beta score of all genes in KO and WT cells. Synthetic lethal hits with MTHFD2 KO with a beta score < -1 are shown in red, and synthetic viable hits with MTHFD2 KO with a beta score > 1 are shown in blue. Shared hits between both KOs are indicated with a black stroke. **i,** Percentage of survivor WT, KO1 and KO2 cells normalized to DMSO after etoposide treatment with indicated concentrations for 72 hours; means + s.d. (*n*=3), unpaired two-tailed *t*-test (*, *P*<0.05; **, *P*<0.01). Inside the graph, a representative scanned image of one replicate (4 technical replicates are shown per condition).

In summary, the analysis of the nuclear interactome of MTHFD2 locates this enzyme in the mitotic environment, where it interacts with several proteins involved in the spindle assembly checkpoint, chromosome segregation and methylation of centromeric DNA and proteins. This suggests a key function of the metabolic enzyme MTHFD2 in mitosis regulation, which is supported by the co-expression analyses and its nuclear accumulation during the G2-M phase transition.

### Centromeric DNA is overexpressed in the absence of MTHFD2

To explore the consequences of MTHFD2 loss, we generated MTHFD2 knock-out (KO) HCT116 cells by CRISPR-Cas9. Two MTHFD2 KO cell lines were validated by sequencing (Extended Data Fig. 5a), Western blot (Fig. 3a) and immunofluorescence (Fig. 3b). Both KO cell lines showed decreased proliferation rates (Extended Data Fig. 5b), invasion (Extended Data Fig. 5c) and clonogenic (Extended Data Fig. 5d) capacities, as previously reported^16, 18, 20^.

**Fig. 5:**
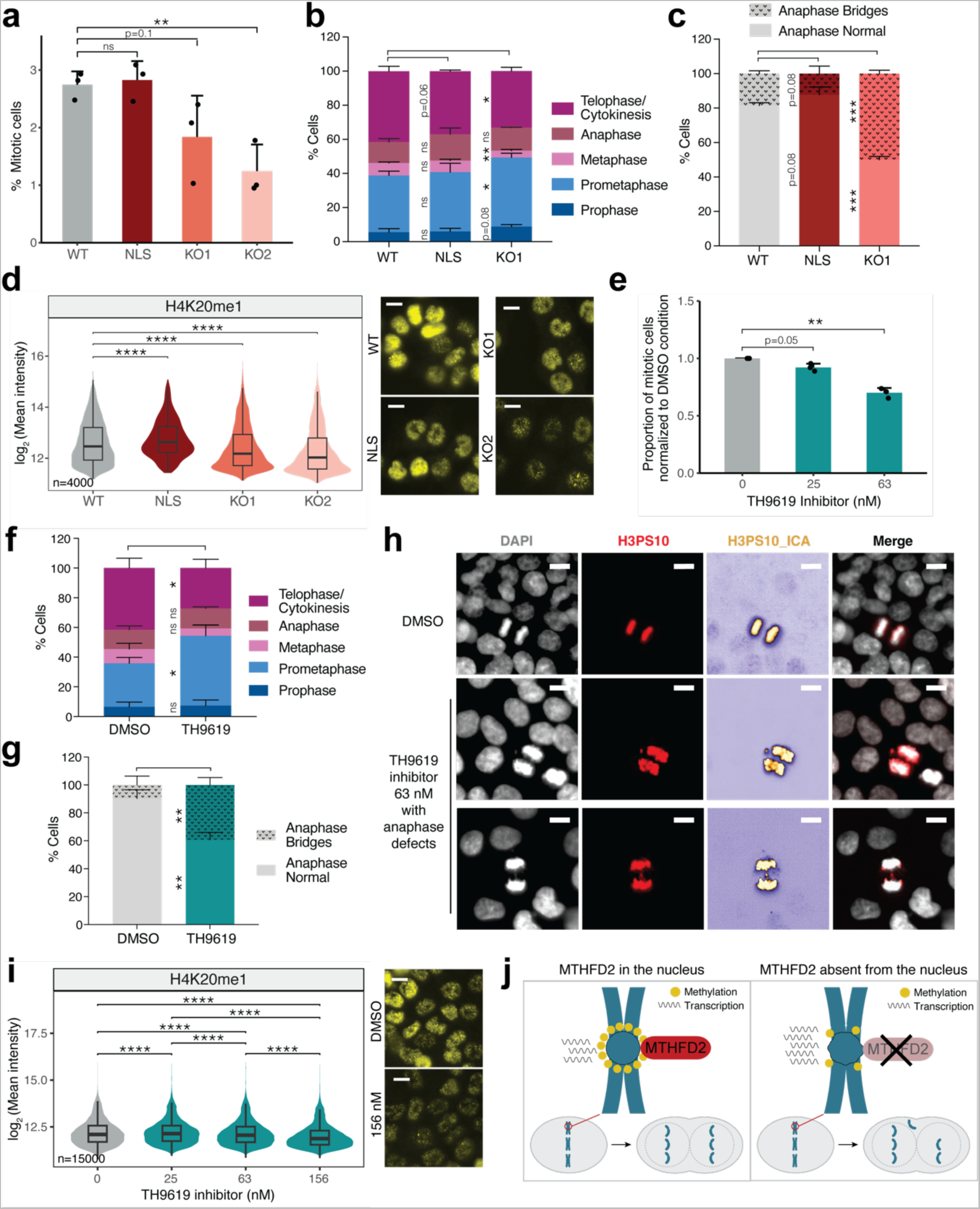
Nuclear MTHFD2 catalytic activity is required for mitosis progression. **a**, Percentage of mitotic cells in HCT116 MTHFD2 wild-type (WT), nuclear (NLS), and knock-out (KO1, KO2) conditions; means + s.d. (*n*=3), unpaired two-tailed *t*-test (ns, non-significant; **, *P*<0.01). **b,** Percentage of WT, NLS and KO1 cells at different mitotic phases; means + s.d. (*n*=3), a minimum of 100 mitotic cells per replicate were analyzed, unpaired two-tailed *t*-test (ns, non-significant; *, *P*<0.05; **, *P*<0.01). **c,** Quantification of anaphase bridges in WT, NLS and KO1 cells; means + s.d. (*n*=3), a minimum of 10 anaphases per replicate were analyzed, unpaired two-tailed *t*-test (***, *P*<0.001). **d,** Comparison of the log_2_ mean intensity of nuclear levels of H4K20me1 in WT, NLS, KO1 and KO2 cells; unpaired two-tailed Wilcoxon test (****, *P*<0.0001). Representative images are shown on the right; non-confocal mode, scale bar 10 μm. **e,** Proportion of HCT116 mitotic cells treated with the indicated concentrations of TH9619 inhibitor for 96 hours normalized to DMSO condition; means + s.d. (*n*=3), one-sample two-tailed *t*-test (**, *P*<0.01). **f,** Percentage of HCT116 cells treated with DMSO or 63 nM TH9619 inhibitor for 96 hours at different mitotic phases; means + s.d. (*n*=3), a minimum of 300 mitotic cells per replicate were analyzed, unpaired two-tailed *t*-test (ns, non-significant; *, *P*<0.05). **g,** Quantification of anaphase bridges in HCT116 cells treated with DMSO or 63 nM TH9619 inhibitor for 96 hours; means + s.d. (*n*=3), a minimum of 40 anaphase cells per replicate were analyzed, unpaired two-tailed *t*-test (**, *P*<0.01). **h,** Representative images of anaphase defects in HCT116 cells treated with DMSO or 63 nM TH9619 inhibitor for 96 hours. DAPI is shown in grey and H3 phospho-Ser10 (H3PS10) is shown in red (second column) or ICA (third column); non-confocal mode, scale bar 10 μm. **i,** Comparison of the log_2_ mean intensity of nuclear levels of H4K20me1 in HCT116 cells treated with DMSO or the indicated concentrations of TH9619 inhibitor for 96 hours; unpaired two-tailed Wilcoxon test (****, *P*<0.0001). Representative images are shown on the right; non-confocal mode, scale bar 10 μm. **j,** Schematic representation of the function of MTHFD2 in the nucleus.

We performed transcriptomics analysis to investigate whether the loss of MTHFD2 could transcriptionally impair cell cycle progression. RNA-sequencing was performed at an early time point (right after KO isolation) and at a late time point (after two months of culturing), to see whether the transcriptome was stable following MTHFD2 KO. We compared HTC116 MTHFD2 wild-type (WT) cells with MTHFD2 KO cells at both time points. The Principal Component Analysis (PCA) indicated that the major difference was the absence of MTHFD2, while time did not influence the transcriptome (Extended Data Fig. 5e), suggesting a rapid and stable transcriptional rewiring of the MTHFD2 KO cells. At the early time point, a total number of 213 genes were differentially expressed between MTHFD2 KO and WT cells, 68 of them upregulated and 145 downregulated after MTHFD2 loss. For the late time point, a total number of 239 genes were differentially expressed, with 98 of them upregulated and 141 downregulated upon MTHFD2 KO (Extended Data Fig. 5f, Supplementary Table 5). From both time points, around 60% of the differentially expressed genes were shared, confirming a good similarity between these samples. Given that MTHFD2 is co-expressed and localized with mitotic proteins, we queried whether the transcriptional rewiring driven by its absence could impact the expression of mitotic genes. Hence, we performed an over-representation analysis of GO Biological Processes with the list of shared up- and downregulated genes upon MTHFD2 KO in both time points. As for downregulated genes, we could retrieve many terms related to metastasis, which agrees with the inferior invasion capability of MTHFD2 KO cells; while for the upregulated genes, we found many terms associated with metabolic processes involving aminoglycans or fatty acids, which suggests a metabolic rewiring following MTHFD2 loss (Extended Data Fig. 5g). We did not retrieve, however, any terms related to mitosis or cell cycle. Likewise, Gene Set Enrichment Analysis (GSEA) did not identify any gene sets associated with mitosis or cell cycle, although we observed a negative enrichment of the PI3K-Akt-mTOR signaling in the KO condition (Extended Data Fig. 5h), which has been previously reported^60^, validating the quality of our dataset.

Since the MTHFD2 pull-down revealed that MTHFD2 is associated with proteins involved in centromeric functions, we assessed the expression of centromeres after MTHFD2 KO. Centromeres consist of alpha-satellite repeats which are transcribed into non-coding RNA that enable the centromeric recruitment of the histone H3 variants CENP-A and CENP-C^42^. As a consequence, an altered centromeric transcription results in defective centromere and kinetochore assembly^42, 61–63^. To assess centromeric expression, we calculated the percentage of reads that intersected with centromeric regions normalized by the total number of mapped reads. We observed that the percentage of reads intersecting with the centromeres was significantly higher in the KO condition compared to the WT counterpart (Fig. 3c), indicating a dysregulation. To validate this observation, we performed retro-transcription quantitative PCR of 20 chromosomal centromeres, which showed that centromeric expression was generally upregulated in the absence of MTHFD2 in all the chromosomes, although only significantly in chromosomes 11 and 20 (Fig. 3d), probably due to the high variability of expression of these regions which may depend on cell cycle^64^. We also inspected the expression of a list of proteins involved in the centromere-kinetochore structure^65^ and we did not observe differential expression following MTHFD2 loss. Of note, only ZWINT and CENP-C were respectively significantly up- and downregulated upon MTHFD2 KO (Extended Data Fig. 5i).

Overall, these results show that the transcriptional switch induced by MTHFD2 loss does not directly impact mitosis or the cell cycle. However, centromeres are aberrantly overexpressed in the absence of MTHFD2, which may lead to centromeric instability and mitotic defects.

### Loss of MTHFD2 decreases centromeric histone and DNA methylation

Upregulation of centromeric expression may arise from an aberrant relaxation of centromeric chromatin upon MTHFD2 loss. Given the fact that our interactome analysis revealed MTHFD2 nuclear interactions with centromeric histone and DNA methyltransferases (KMT5A and DNMT3B), we queried whether the absence of MTHFD2 could impair centromere methylation. We observed a significant decrease of nuclear H4K20me1 levels, a specific centromeric histone mark deposited by KMT5A^33, 43, 56^, upon MTHFD2 absence, with minor changes in H3K9me3 and an overall mild increase in H3K27me3 (Fig. 3e). Besides, we measured the coefficient of variation of the three histone marks, which assesses the nuclear heterogeneity of the staining reflecting the compartmentalization into eu- and heterochromatin. Only H4K20me1 showed a clear deviation towards less variation in the absence of MTHFD2 (Extended Data Fig. 5j). We then quantified the levels of H4K20me1, together with H3K9me3 and H3K27m3, at centromeric regions using chromatin immunoprecipitation (ChIP) coupled to quantitative PCR. We observed that, at centromeres, all these histone marks were considerably reduced (Fig. 3f-h), fitting the hypothesis of a centromeric transcriptional upregulation following a demethylation-induced chromatin relaxation in the absence of MTHFD2. DAPI staining, however, also suggested a global chromatin reorganization upon MTHFD2 loss, since the coefficient of variation of DAPI decreased in the absence of MTHFD2, pointing to a more homogeneous DNA distribution (Extended Data Fig. 5k). Given the fact that we observed an MTHFD2 interaction with DNMT3B, we investigated whether the loss of DNA methylation could explain the observed changes in DAPI staining upon MTHFD2 KO. We performed Nanopore whole-genome sequencing and conducted a comparative methylation analysis of HTC116 MTHFD2 WT and KO cells (Extended Data Fig. 6a-d, Supplementary Table 6). We categorized the differentially methylated sites into hypermethylated or hypomethylated if they were significantly more or less methylated, respectively, in the MTHFD2 KO condition compared to the WT. We observed a strong hypomethylation in MTHFD2 KO cells either when looking at CpG sites located at CpG islands and shores (Fig. 3i) or considering all CpG sites (Extended Data Fig. 6e), being the centromeres equally affected as the rest of the genome.

Annotation of the genomic location of the hyper- and hypomethylated sites showed that they largely fell in intergenic regions (61% and 65%, respectively) (Extended Data Fig. 6f,g), which may suggest that the observed methylation changes should not directly affect the transcriptome of the MTHFD2 KO cells. Indeed, differentially methylated regions were equally distributed into up- and downregulated genes (Extended Data Fig. 6h). Likewise, differentially expressed genes were also equally distributed into hyper- and hypomethylated regions (Extended Data Fig. 6i), reinforcing the lack of correlation between changes in DNA methylation and gene expression following MTHFD2 loss.

Our data indicate that MTHFD2 KO cells present a strong methylation defect at the level of both histones, particularly at the centromere, and DNA, mostly present in non-coding regions, which may affect genome stability but does not impact gene expression.

### In the absence of MTHFD2, cells accumulate chromosomal alterations

Following our previous observation of strongly hypomethylated intergenic regions in MTHFD2 KO cells, we hypothesized that this lack of methylation could affect chromatin stability and result in chromosome alterations. Thus, we karyotyped the HTC116 MTHFD2 WT and KO cells to see whether they have acquired any chromosomal alteration. HTC116 cells are well-suitable for this type of analysis since they are near diploid. We observed that the karyotype was mostly the same in the three cell lines, as expected, although both KO cell lines each presented an additional alteration that was not displayed in the WT population (Extended Data Fig. 6j-l). The MTHFD2 KO1 had a deletion in the small arm of chromosome 6 (del(6)(p12)), while the MTHFD2 KO2 showed an insertion in the small arm of chromosome 14 (add(14)(p11)) (Fig 3j, Extended Data Fig. 6j-l). Interestingly, both alterations were located very close to the centromeric region, suggesting that they may have arisen due to centromeric instability.

Given the limitation of the karyotype technique in identifying smaller genomic alterations, we queried the Nanopore whole-genome sequencing data presented above to investigate whether the loss of MTHFD2 was associated with an accumulation of genomic structural variation. Although MTHFD2 WT and KO cells shared a considerable number of genetic variants (Extended Data Fig. 6m), MTHFD2 KO cells diverged from the WT population by showing a higher number of each type of variant (Fig. 3k). The size distribution of the variants, which could span from a few bases to 100 kb, was similar in both conditions (Extended Data Fig. 6n). We excluded that deletions could be the reason of the observed hypomethylation by checking the proportion of hypomethylated and hypermethylated sites located at deleted regions, which did not change across conditions (Supplementary Table 7). Finally, we performed a centromere-focused analysis and observed that also there the absence of MTHFD2 induced a higher number of alterations (Fig. 3l).

Taken together, our data show that MTHFD2 loss leads to genomic aberrations, which may be the result of mitotic defects arising from centromeric and genomic instability.

### MTHFD2 is required for proper chromosome segregation

Our pull-down MS and co-expression analyses indicated that nuclear MTHFD2 plays a role in mitosis. Besides, we have identified that, in the absence of MTHFD2, centromeric expression is aberrantly increased, following centromeric histone and DNA demethylation defects.

We next explored whether MTHFD2 loss leads to mitotic errors, which may link the centromere expression and methylation defects with the accumulation of genomic variants. First, we evaluated the mitotic index, that is the percentage of cells in mitosis, in MTHFD2 WT and KO HTC116 cells, measuring the mitotic marker histone H3 phospho-Ser10 (H3PS10) by high-throughput immunofluorescence. The mitotic index was reduced upon MTHFD2 KO (Fig. 4a and Extended Data Fig. 7a). In agreement, we observed that the number of G2/M cells was decreased in both KO cell lines (Extended Data Fig. 7b). We then asked whether MTHFD2 WT and KO cells needed the same time to complete mitosis. We synchronized the cells at the G2-M border by treating them with the CDK1 inhibitor RO-3306 for 20 hours. Then, we released them and assessed the percentage of cells in G1 at different time points. While MTHFD2 WT cells quickly proceeded through mitosis with ∼70% of cells back in G1 after 2.5 hours, both MTHFD2 KO cell lines showed a mitotic delay and only ∼50% of cells were in G1 at the same time point (Fig. 4b and Extended Data Fig. 7c). We reasoned that the delay in mitosis progression could be due to a defect in one of the phases of mitosis. Classifying mitotic cells into the different phases showed an imbalance in their proportion in comparison to the WT counterpart. Overall, MTHFD2 absence retained cells in early mitotic phases (prophase and prometaphase) for a longer time (Fig. 4c), suggesting a defect of chromosome congression at the metaphase plate. MTHFD2 loss also resulted in a decreased proportion of cells in telophase and cytokinesis (Fig. 4c), which could arise from the inability to repair chromosome segregation defects in anaphase.

To validate this hypothesis, we analyzed metaphases in both conditions and discovered that the loss of MTHFD2 increased the percentage of metaphase plates lagging one chromosome as shown by the anti-kinetochore (CREST) staining, confirming the chromosome congression defect (Fig. 4d,e). Defects in chromosome congression may result in impaired chromosome segregation which, ultimately, leads to genomic aberrations as the ones observed in our MTHFD2 KO cells. We therefore investigated whether the absence of MTHFD2 induced chromosome segregation defects in anaphase. MTHFD2 KO cells indeed displayed a higher number of DNA anaphase bridges (Extended Data Fig. 7d,e), clearly manifesting difficulties in segregating chromosomes. We thus performed CREST staining to determine whether the observed anaphase bridges involved the centromere. In the MTHFD2 KO cells, more than 21% of the anaphases displayed defects involving the centromeres (Fig. 4f,g), corroborating the hypothesis of an MTHFD2 loss-induced centromeric instability. Lagging chromosomes or chromosomal fragments that contain centromeric regions usually give rise to CREST-positive micronuclei. Analysis of interphase cells showed that the number of CREST-positive micronuclei was significantly increased in both MTHFD2 KO cell lines as compared to the WT parental population (Extended Data Fig. 7f,g).

Since nuclear MTHFD2 interacts with key regulators of the spindle assembly checkpoint (TPR, MAD1L1), we observed a reduced percentage of mitotic cells in metaphase in both KO conditions (Fig. 4c) and defects in chromosome congression (Fig. 4d,e), we queried whether MTHFD2 could have a role in the activity of the spindle assembly checkpoint. We performed a correlation analysis between MTHFD2 RNA expression and key genes involved in the spindle assembly checkpoint^65^ (AURB, BUB1, BUB1B, BUB3, CCNB1, CDC20, MAD1L1, MAD2L1, MD2BP, TKK, ZW10 and ZWILC) using TCGA patient expression data. Indeed, MTHFD2 was strongly positively correlated with 11 of these 12 genes (Extended Data Fig. 8a-l). Additionally, since MTHFD2 KO cells exhibited more mitotic defects, we asked whether MTHFD2 expression levels correlate with cell ploidy. For this, we retrieved the aneuploidy status and score from all the cancer cell lines in the CCLE database^47, 66^, along with MTHFD2 expression and essentiality values. We detected an inverse association between MTHFD2 expression and aneuploidy class, where near-euploid cell lines had the highest MTHFD2 expression, while highly-aneuploid cell lines had the lowest MTHFD2 expression (Extended Data Fig. 8m). In agreement, cells with high MTHFD2 expression (top 25%) showed a decreased aneuploidy score compared to low MTHFD2 expression (bottom 25%) cells (Extended Data Fig. 8n). We then explored whether there was a relationship between MTHFD2 CRISPR essentiality and aneuploidy status in cells with high MTHFD2 expression (top 25%) and we observed an inverse correlation, with near-euploid cells having the highest MTHFD2 essentiality while high-aneuploid cells the lowest (Extended Data Fig. 8o). The fact that near-euploid cells have higher MTHFD2 expression and essentiality further supports the hypothesis that MTHFD2 is needed for correct chromosome segregation in mitosis.

Next, we undertook an orthogonal and unbiased approach to verify that MTHFD2 loss impacts mitotic progression. We reasoned that a CRISPR-Cas9 genetic screen on the MTHFD2 KO cells should identify as synthetic lethal genes that favor mitosis progression, while as synthetic viable genes that slow it down. We performed a genome-wide CRISPR-Cas9 genetic screen by transducing MTHFD2 WT and KO cells. Cells were selected with puromycin for 8 days, where the initial population was harvested, and then kept in culture for three more weeks, where the final population was harvested. For the three samples, DNA from the initial and final population was extracted and sequenced. The average coverage of the initial population for all the samples was higher than 500X (Extended Data Fig. 9a-c), and the Gini index was lower than 0.1 in all initial samples (Extended Data Fig. 9d), ensuring the evenness of sgRNA read counts^67^. A cell-cycle normalization was performed to compensate for cell cycle differences among conditions (Extended Data Fig. 9e-f). Finally, enriched and depleted sgRNAs were identified by comparing each final population with their corresponding initial population, and enriched and depleted sgRNAs in each KO cell line were compared with those in the WT to identify synthetic lethal genes or synthetic viable genes with MTHFD2 loss. Genes detected in both KO (beta(KO)-beta(WT) < -1 and > 1) were further considered (Fig. 4h, Supplementary Table 8).

Confirming the role of MTHFD2 in the regulation of the mitotic process, we retrieved key mitotic players in both directions. For instance, the loss of WAPL, which provokes chromosome condensation by locking Cohesin on chromatin^68^, was found to be synthetically viable with MTHFD2 KO (Fig. 4h). On the other hand, TOP2A was found among the top synthetic lethal genes with MTHFD2 KO (Fig. 4h). Opposite to WAPL, TOP2A is required for mitotic chromosome condensation^69^, and given the methylation defects observed in the MTHFD2 KO cells, any additional problems of chromatin condensation may be fatal. Treatment with the Topoisomerase II inhibitor etoposide confirmed the higher sensitivity of MTHFD2 KO cells to the loss of TOP2A (Fig. 4i).

Overall, our results demonstrate that the lack of MTHFD2 induces a delay in chromosome congression, chromosome segregation defects and accumulation of centromere-containing micronuclei, reinforcing the connection between MTHFD2-dependent centromere and chromosome stability and mitosis regulation.

### Nuclear MTHFD2 catalytic activity is required for mitosis progression

We reasoned that, if MTHFD2 controls the methylation of centromeric histones and DNA that is required for proper mitosis progression, the supplementation of S-adenosyl methionine (SAM) or folate derivatives could potentially restore the mitotic index. Thus, we treated MTHFD2 WT and KO cells with two concentrations of formate, folate, 5,10-methylenetetrahydrofolate (MTHFD2 product) or SAM for 72 hours, being the lowest concentration near plasma physiological levels. We did not observe any recovery in the mitotic index in any of the KO cell lines (Extended Data Fig. 10a). This result suggested two scenarios, being the first one a structural role of MTHFD2 required for DNA and histone methylation or, the second, a nuclear MTHFD2 enzymatic function required to keep a certain nuclear metabolic environment. We therefore generated a nuclear-MTHFD2 (NLS) HTC116 cell line by introducing a triple nuclear localization signal in the MTHFD2 locus by CRISPR knock-in (Extended Data Fig. 10b,c). We compared the mitotic index of MTHFD2 NLS cells to those of either MTHFD2 WT and KO cells and observed that the nuclear restriction of MTHFD2 allowed for a mitotic index comparable to the one of WT cells (Fig. 5a). Additionally, MTHFD2 NLS cells grew at a very similar rate than MTHFD2 WT cells (Extended Data Fig. 10d), recovering the proliferation defect observed in the MTHFD2 KO population (Extended Data Fig. 5b). Similarly, the mitotic phase distribution and the rate of anaphase bridges observed in the MTHFD2 NLS cells were similar to the ones of the MTHFD2 WT cells (Fig. 5b,c). We also assessed global H4K20me1 levels, which indicated that the MTHFD2 NLS cells do not show the histone methylation defect present in the KO condition and had slightly higher levels of H4K20me1 than the MTHFD2 WT cells (Fig. 5d, Extended Data Fig. 10e). In line with this result, in the MTHFD2 NLS cells, the coefficient of variation of DAPI, which was decreased upon MTHFD2 loss (Extended Data Fig. 5k), was not only recovered but higher than in WT cells (Extended Data Fig. 10f).

To rule out whether the nuclear enzymatic function of MTHFD2 is required for mitosis control, we employed a recently published MTHFD2 inhibitor TH9619^23^ that cannot enter the mitochondria^70^, hence behaving as a nuclear MTHFD2 inhibitor. When we treated the cells with non-toxic concentrations of this compound (Extended Data Fig. 10g) for 96 hours, we observed a dose-response decrease of the mitotic index (Fig. 5e). Additionally, the treatment induced an imbalance of the mitotic phases (Fig. 5f), similar to the one observed in the MTHFD2 KO population (Fig. 4c). We also evaluated anaphase defects and observed that the treatment doubled them (Fig. 5g,h). In parallel, we investigated the levels of H4K20me1 and observed a significant decrease in the presence of the treatment (Fig. 5i, Extended Data Fig. 10h), although the coefficient of variation of DAPI did not noticeably change (Extended Data Fig. 10i). These results indicate that, while nuclear MTHFD2 enzymatic activity is required for the correct distribution of mitotic phases and to achieve successful chromosome segregation, mitochondrial MTHFD2 seems to be irrelevant for this process.

Overall, our results show that the non-canonical localization of MTHFD2 in the nucleus is required for proper DNA and centromeric histone methylation and centromere transcription.

Perturbation of MTHFD2 nuclear function leads to chromosomal instability, incorrect mitosis progression and accumulation of genetic aberrations (Fig. 5j).

## Discussion

Nuclear metabolism is an emerging field of biology^1–5^. The evidence that metabolic reactions can happen in unexpected cellular compartments, such as the nucleus, is revolutionizing the classical idea of central metabolism and suggests an on-demand production of metabolites to satisfy precise cellular needs. Clear examples of nuclear metabolism events are the production of nuclear acetyl-CoA^6, 8, 9^, ATP^71^, SAM^7^ and nucleotides^26^.

In the last decade, the folate enzyme MTHFD2 has become a promising therapeutic target, since it is upregulated and drives cancer progression in a wide range of cancer types^13– 16, 18–20^. Consequently, several MTHFD2 inhibitors have been already developed with favorable pre-clinical results^21–23^. The canonical function of MTHFD2 appears to contribute to tumorigenesis by providing substrates for *de novo* synthesis of purines and pyrimidines^72^, or by maintaining the redox state of the cell^15, 73^. However, MTHFD2 does not perform an exclusive enzymatic reaction within the folate pathway and its catalytic activities are shared with other members of the MTHFD family^10^. Therefore, it is not clear why the cancer cell increases the expression of MTHFD2.

Recent publications have shown that MTHFD2 can localize within the nuclear environment^24–26, 74^, where it might regulate RNA translation and metabolism^75^, DNA damage repair^74^ or even bind to chromatin^26^. However, a detailed understanding of the cellular processes that determine the requirement for MTHFD2 in the nucleus has not yet been achieved, nor whether its enzymatic function there is necessary. Here we demonstrated for the first time that enzymatically active MTHFD2 is recruited to the nucleus during the G2-M phase of the cell cycle, and is required to maintain methylation of centromeres, thus preventing their transcriptional upregulation. Genetic or pharmacological disruption of this safeguard mechanism leads to aberrant mitosis, resulting in chromosomal instability that compromises cancer cell survival.

After proving MTHFD2 nuclear localization across a wide variety of cell lines (Fig. 1b,c) and patient-derived colon organoids (Extended Data Fig. 1h), we characterized the MTHFD2 nuclear interactome by coupling the pull-down of the enzyme following subcellular fractionation to mass spectrometry. We noticed that a considerable number of nuclear proteins were detected as MTHFD2 interactors in the cytosolic fraction (Extended Data Fig. 2c). Although this situation could arise from technical issues with the fractionation protocol, another plausible cause could be mitotic cells. During the first phases of mitosis, the nuclear envelope breakdown exposes the nuclear content to the cytoplasmic fraction, thus provoking a natural nuclear contamination of the cytosolic compartment. Corroborating this latter hypothesis, most of the nuclear MTHFD2 interactors retrieved in the cytosolic compartment were mitotic players (Supplementary Table 2). Therefore, we called nuclear MTHFD2 interactors those proteins with an annotated nuclear localization either retrieved in the chromatin or the cytoplasm MTHFD2 pull-downs (Fig. 2a, Extended Data Fig. 2c).

Classification of the nuclear MTHFD2 interactors into functional categories revealed a possible role for nuclear MTHFD2 regulating mitosis progression (Fig. 2a). Transcriptomics and proteomics co-expression analyses (Fig. 2c, Extended Data Fig. 3f) not only supported the connection between MTHFD2 and mitosis but also pointed towards specific mitotic processes involving chromosome segregation in anaphase. Interestingly, a potential cell cycle-related role of MTHFD2 has been previously suggested^25, 76^, although never fully addressed. The accumulation of nuclear MTHFD2 in the G2-M phase of the cell cycle supports its mitotic role (Fig. 2e,g). Even though in the present study we do not investigate how MTHFD2 translocates into the nucleus, we hypothesize that this phenomenon may be allowed by the cell cycle-dependent mitochondria fission that happens early in mitosis^77^, which would further support the observed spatiotemporal compartmentalization of the enzyme.

The transcriptomics analysis showed that MTHFD2 loss provokes an aberrant overexpression of the centromeric regions (Fig. 3c,d). Alterations of centromere expression are associated with kinetochore malfunctioning and mitotic aberrations^42, 61–63^. We reasoned that centromere overexpression may be a result of chromatin decompaction following MTHFD2 KO. The methyltransferase KMT5A is a nuclear MTHFD2 interactor (Fig. 2a) and the writer of H4K20me1^43^, whose deposition at the CENP-A centromeric nucleosome is essential for kinetochore assembly^33, 43, 56^. Nuclear levels of H4K20me1 significantly decreased in the absence of MTHFD2 (Fig. 3e). When specifically checking the centromeric regions, we observed that the methylation defect goes beyond H4K20me1 and affects also the levels of the repressive marks H3K9me3 and H3K27me3 (Fig. 3f-h), supporting the hypothesis of a chromatin demethylation-mediated centromeric upregulation.

In the transcriptomic analysis, we did not identify a clear mitotic gene signature in the absence of MTHFD2, indicating that the involvement of MTHFD2 in mitosis is not gene expression-related but rather through a more direct mechanism. Of the centromere and kinetochore-associated proteins, only ZWINT and CENP-C were significantly up- and downregulated, respectively (Extended Data Fig. 5i). Of interest, CENP-C loss has been previously reported to increase alpha-satellite transcription^78^, which could either synergize with the MTHFD2 loss-driven centromeric hypomethylation or be a consequence of it.

DNA methylation was also largely reduced in the absence of MTHFD2 (Fig. 3i, Extended Data Fig. 6e). DNA hypomethylation has been widely linked to chromosomal instability^37–40^. Interestingly, the loss of function of DNMT3B, a nuclear interactor of MTHFD2 (Fig. 2a), drives chromosomal instability through DNA hypomethylation^79, 80^.

A karyotype analysis showed that MTHFD2 KO cells had pericentromeric chromosomal defects (Fig. 3j, Extended Data Fig. 6j-l), and whole-genome Nanopore sequencing revealed that they accumulate more structural variants than their WT counterpart (Fig. 3k). Structural variants are fixed in the DNA during DNA replication, being cell division-dependent. MTHFD2 KO cells showed reduced cell proliferation (Extended Data Fig. 5b) when compared to the WT cells. Hence, correcting for the proliferation rate of each cell population may even increase the divergence in structural variation accumulation observed between the MTHFD2 WT and KO cells.

Supporting the role of MTHFD2 in centromere and chromosomal stability, in the absence of MTHFD2, we observed concrete mitotic defects, such as defects in chromosome congression (Fig. 4d,e) and segregation (Fig. 4f,g, Extended Data Fig. 7d,e), along with an increased number of centromere-containing micronuclei (Extended Data Fig. 7f,g). With an orthogonal functional genomics approach, we discovered that the mitotic players WAPL and TOP2A are, respectively, synthetic viable and synthetic lethal with MTHFD2 KO (Fig. 4h).

WAPL depletion induces chromatin condensation by locking Cohesin on the DNA^68^. Thus, the absence of WAPL could compensate for the MTHFD2 loss-driven chromatin decompaction caused by DNA and histone methylation defects. On the contrary, TOP2A loss induces chromosome structural defects and mitotic delay^69^, which would exacerbate the hypomethylation and chromosomal instability phenotype observed in the absence of MTHFD2, explaining the synthetic lethality.

To show that the nuclear pool of MTHFD2 is crucial for the proper progression of mitosis, we demonstrated that nuclear restriction of MTHFD2 does not impair the mitotic index (Fig. 5a), nor does it reduce H4K20me1 (Fig. 5d) or exhibit anaphase defects (Fig. 5c). Interestingly, these results suggest that the mitochondrial function of MTHFD2 is dispensable for correct mitosis progression. Furthermore, nuclear-restricted MTHFD2-expressing cells do not show an obvious proliferative defect compared with WT cells (Extended Data Fig. 10d), casting doubts on whether the pivotal role of MTHFD2 in cancer may be related to its canonical mitochondrial function.

To test whether the enzymatic activity of MTHFD2 is required in the nucleus, we used a recently published MTHFD2 inhibitor^23^ that cannot pass the mitochondrial membrane, thus behaving as a nuclear-MTHFD2 inhibitor^70^. The compound can also inhibit MTHFD1, the cytoplasmic MTHFD2 homolog, and induces cell death by provoking an imbalance of cytoplasmic folate derivatives^70^. When cells were cultured in the presence of thymidine, which counteracts the effect mediated by MTHFD1 inhibition, even at subtoxic concentrations of the compound, we observed a reduction in the cellular mitotic index (Fig. 5e), an increase in anaphase defects (Fig. 5f,g) and a decrease in H4K20me1 levels (Fig. 5h), recapitulating the phenotype triggered by the loss of MTHFD2.

Intriguingly, the product of MTHFD2, 5,10-methylenetetrahydrofolate, *per se* cannot directly contribute to DNA or histone methylation, which requires SAM. 5,10-methylenetetrahydrofolate needs to be converted into SAM by two sequential catalytic reactions performed by Methylenetetrahydrofolate Reductase (MTHFR) and Methionine Synthase (MTR)^10^, which were not found among our MTHFD2 interactors, nor on chromatin^26^. This apparent controversy suggests that the nuclear enzymatic function of MTHFD2 may be required to keep a suitable metabolic environment that ensures adequate centromeric methylation and stability, rather than to methylate centromeres directly. Additionally, the treatment with MTHFD2 downstream metabolites did not rescue the mitotic index defect observed in MTHFD2 KO cells (Extended Data Fig. 10a). These results suggest that MTHFD2 might fulfill a precise local metabolic requirement that cannot be fully met by increasing absolute metabolite levels, underlying the fundamental importance of nuclear metabolism compartmentalization.

## Methods

### Cell culture

A459 (ATCC; #CCL-185), BT-549 (ATCC; #HTB-122), EBC-1 (Cellosaurus; #CVCL_2891), HCT 116 (ATCC; #CCL-247), HEK293T (ATCC; #CRL-3216), HT-29 (ATCC; #HTB-38), MCF7 (ATCC; #HTB-22), MDA-MB-231 (ATCC; #HTB-26), RKO (ATCC; #CRL-2577), SK-BR-3 (ATCC; #HTB-30), SW480 (ATCC; #CCL-228), SW620 (ATCC; #CCL-227), T-47D (ATCC; #HTB-133) and U-2 OS (ATCC; #HTB-96) cells were cultured in DMEM (Gibco; #11966025) supplemented with 10% fetal bovine serum (FBS) (Gibco; #10270106) and 1% penicillin/streptomycin (Gibco; #15140122) at 37°C in 5% CO_2_. H1437 (ATCC; #CRL-5872), H226 (ATCC; #CRL-5826) and H358 (ATCC; #CRL-5807) cells were cultured in RPMI GlutaMAX (Gibco; #61870036) supplemented with 10% FBS (Gibco; #10270106) and 1% penicillin/streptomycin (Gibco; #15140122) at 37°C in 5% CO_2_. Cell cultures were tested every month for mycoplasma contamination.

### Plasmids and primers

The plasmids and primers (including gene blocks) used in this study are listed in Supplementary Methods Table 1 and 2, respectively.

To generate the plasmids needed for CRISPR-Cas9 KO/knock-in, Brand and Winter protocol was followed^81^. Briefly, to obtain the cutting vectors sgMTHFD2ex4_GW223 and sgMTHFD2int1-2_GW223, primers with sense and antisense sgRNA sequences (primers 1-4) were designed to generate the sgRNA. The cutting vector GW223_pX330A_sgX_sgPITCh (2 μg) was digested with BbsI-HF (New England Biolabs; #R3539) in Cutsmart Buffer (New England Biolabs; #B6004) for 1 hour, dephosphorylated with Shrimp Alkaline Phosphatase (rSAP) (New England Biolabs; #M0371) and gel purified with the QIAquick PCR & Gel Cleanup Kit (Qiagen; #28506). Sense and antisense oligos were annealed with T4 Polynucleotide Kinase (PNK) (New England Biolabs; #M0201) and ligated with the digested cutting plasmid with T4 DNA ligase (New England Biolabs; #M0202). The ligated fragments were transformed into DH5α E. coli competent cells (Thermo Fisher Scientific; #18265017) and single colonies were analyzed with Sanger sequencing (Eurofins) to select positive clones (primer 5).

To obtain the repair vector 3xNLS-dTAG-GFP-NEO_GW209 several steps were followed. To obtain first the 1xNLS-dTAG-GFP_GW209 plasmid, the repair vector GW209_pCRIS-PITChv2-C-dTAG-Puro (BRD4) (2 μg) was digested with MluI-HF (New England Biolabs; #R3198) in Cutsmart Buffer for 1 hour, dephosphorylated with rSAP and gel purified with the QIAquick PCR & Gel Cleanup Kit. A gene block (Integrated DNA Technologies) with the repair sequence was designed (dTAG_GFP_NLS_MTHFD2) and cloned into the digested repair vector using the Gibson reaction approach for 2 hours at 50°C, followed by DH5α E. coli cells transformation. Single clones were Sanger sequenced (primers 6-8). This plasmid 1xNLS-dTAG-GFP_GW209 was used as a template to amplify the sequence of FKBP-V-GFP-NLS (primers 9-10) and the plasmid pcDNA3-hLOXL2 was used to amplify the SV40Pr-Neo-term sequence (primers 11-12) using the Phusion High-Fidelity DNA Polymerase (Thermo Fisher Scientific; #F530). These two PCR products were cloned into the digested repair vector GW209_pCRIS-PITChv2-C-dTAG-Puro (BRD4) using the Gibson reaction approach for 2 hours at 50°C, followed by DH5α E. coli cells transformation, generating the vector 1xNLS-dTAG-GFP-NEO_GW209. To introduce a 3xNLS sequence, the vector 1xNLS-dTAG-GFP-NEO_GW209 was sequentially cut with AarI (Thermo Fisher Scientific; #ER1581) and BsmBI (New England Biolabs; #R0580) and gel purified. The obtained backbone was ligated via custom Gibson reaction with the gene block 3xNLS (Integrated DNA Technologies) for 2 hours at 50°C. Afterward, the product was transformed in DH5α competent cells to obtain 3xNLS-dTAG-GFP-NEO_GW209. Single clones were Sanger sequenced (primers 6-7, 13-15).

### Chromatome fractionation

1^10^7^ cells were first lysed in CHAPS (3-cholamidopropyl dimethylammonium 1-propane sulfonate) Buffer (0.5-10% CHAPS in PBS) for 15 minutes to break the cytosolic membrane and centrifuged for 5 min at 720g at 4°C. The concentration of CHAPS was optimized for each cell line. The supernatant was harvested as the cytosolic fraction. The nuclear pellet was resuspended in Cytoplasmic Lysis Buffer (IGEPAL 0.1%, NaCl 150 mM, Tris-HCl 10 mM pH 7 in H_2_O), placed on the top of a Sucrose Gradient Buffer (NaCl 150 mM, sucrose 25%, Tris-HCl 10 mM pH 7 in H_2_O) and centrifuged for 5 min at 1200g at 4°C. Purified nuclei were then washed 3 times by resuspending in Nuclei Washing Buffer (EDTA 1 mM, IGEPAL 0.1% in PBS) and centrifuged for 5 min at 1200g at 4°C. Then, the washed nuclear pellet was resuspended in Nuclei Resuspension Buffer (EDTA 1 mM, NaCl 75 mM, 50% sucrose, Tris-HCl 20 mM pH 8 in H_2_O) and the nuclear membrane was lysed by adding Nuclei Lysis Buffer (EDTA 0.2 mM, HEPES 20 mM pH 7.5, IGEPAL 0.1%, NaCl 300 mM in H_2_O), vortexing and incubating for 5 min. After centrifugation for 2 min at 16000g at 4°C, the resulting chromatin was resuspended in Benzonase Digestion Buffer (15 mM HEPES pH 7.5, 0.1% IGEPAL, TPCK 5 μg/mL) and sonicated on a Bioruptor Pico (Diagenode) for 15 cycles 30 sec ON/30 sec OFF in 1.5 mL Diagenode tubes (Diagenode; #C30010016). Finally, sonicated chromatin was digested with benzonase enzyme (VWR; #706643; 2.5U) for 30 min at room temperature, and the resulting sample was harvested as chromatome fraction. All the steps were performed on ice and all buffers were supplemented with proteinase inhibitors (Roche; #4693132001). Cytosolic and chromatome extracts were quantified with Pierce BCA Protein Assay Kit (Thermo Scientific; #PIER23225).

### SDS-electrophoresis and Western blot

Samples were mixed with 4X Laemmli sample buffer (Bio-Rad; #1610747) and boiled at 95°C for 5 min. Proteins were separated by sodium dodecyl-sulfate (SDS)–polyacrylamide gel electrophoresis, and transferred to a nitrocellulose membrane by wet transfer (10% Transfer buffer, 20% methanol in H_2_O). Membranes were blocked in 5% milk (Millipore; #70166) in 0.05% Tween20 in PBS for 1 hour at room temperature. Membranes were incubated with primary antibodies prepared in 0.05% Tween20 in PBS overnight at 4°C. Fluorescent secondary antibodies rabbit-800 (Thermo Fisher Scientific; #A32735; 1:10000) and mouse-780 (Thermo Fisher Scientific; #A21058; 1:10000) were also prepared in 0.05% Tween20 in PBS and incubated for 1 hour at room temperature. Three washes after the primary and secondary antibodies were performed with 0.05% Tween20 in PBS. Detection was achieved with Odyssey CLx (Li-Cor) and analyzed with Image Studio Lite (version 5.2.5).

For the chromatome experiments, the following primary antibodies were used: MTHFD2 (Abcam; #ab151447; 1:1000), Vinculin (Cell Signaling; #13901S; 1:1000), FDX1 (Thermo Fisher Scientific; #PA559653; 1:1000) and H3 (Cell Signaling; #14269S; 1:1000).

For pull-down analysis, the following primary antibodies were used: MTHFD2 (Abcam; #ab151447; 1:1000), MTHFD2 (Abcam; #ab56772; 1:700), TPR (Santa Cruz Biotechnology; #sc-101294; 1:1000), KIF4A (Santa Cruz Biotechnology; #sc-365144; 1:1000), PRMT1 (Santa Cruz Biotechnology; #sc-166963; 1:1000) and KMT5A (Thermo Fisher Scientific; #PA5-31467; 1:2000).

For validating the MTHFD2 KO, whole-cell extracts were obtained using an SDS lysis buffer (2% SDS, 50 mM Tris-HCl pH 7, 10% glycerol in H_2_O), and the following primary antibodies were used: MTHFD2 (Abcam; #ab151447; 1:1000) and Vinculin (Cell Signaling; #13901S; 1:1000).

### Immunofluorescence

Immunofluorescence experiments were performed by seeding cells on clear flat-bottom 96-well plates (Perkin Elmer; #6055302) and fixing them with 4% formaldehyde (Thermo Fisher Scientific; #28908) for 15 min at room temperature. Permeabilization was performed using 0.2% Triton X-100 in PBS for 30 min, followed by blocking with 2% bovine serum albumin (BSA) in PBS for 45 min. Cells were incubated first with primary antibodies for 1 hour at room temperature. The following antibodies were used: MTHFD2 (Abcam; #ab151447; 1:500), H3PS10 (Sigma-Aldrich; #06-570; 1:500), α-tubulin (Sigma-Aldrich; #T9026; 1:500), CREST (AntibodiesInc; #15-234; 1:500), H4K20me1 (Diagenode; #C15410034; 1:500), H3K9me3 (Diagenode; #C15410193; 1:1000) and H3K27me3 (Diagenode; #C15410195; 1:200). After washing, cells were incubated with secondary antibodies for 1 hour at room temperature in the dark. The following antibodies were used: Alexa Fluor 488 donkey anti-rabbit (Thermo Fisher Scientific; #A21206; 1:1000), Alexa Fluor 647 goat anti-rabbit (Thermo Fisher Scientific; #A21244; 1:1000), Alexa Fluor 555 goat anti-mouse (Thermo Fisher Scientific; #A21058; 1:1000) and Alexa Fluor 647 goat anti-human (Thermo Fisher Scientific; #A21445; 1:1000). Finally, cells were incubated with DAPI (4,6-diamidino-2-phenylindole) (Sigma-Aldrich; #MBD0015; 1:1000) for 5 min at room temperature in the dark (except FUCCI cells). After the incubations with antibodies and DAPI, cells were washed twice with 0.05% Tween20 in PBS and once with PBS. Images were taken with the Operetta High Content Screening System (PerkinElmer) using a 10X, 40X or 63X objective and non-confocal or confocal mode.

Images were quantified using the Harmony software (version 4.9), first by identifying nuclei and then quantifying their properties (H3PS10, H4K20me1, H3K9me3, H3K27me3 and DAPI mean intensity and/or coefficient of variation). For calculating the mitotic index, individual cells with a signal higher than 3 standard deviations of the average H3PS10 mean intensity were considered mitotic cells. For the quantification of the epigenetic marks and DAPI, the individual mean intensity and coefficient of variation of H4K20me1, H3K9me3, H3K27me3 and DAPI were considered.

The classification of mitotic cells and the identification and quantification of mitotic defects were performed with ImageJ (version 1.52q) using the DAPI, H3PS10, CREST and α-tubulin staining.

To quantify the percentage of survival of HCT116 cells after TH9619 treatment, cells were fixed and permeabilized as previously indicated and directly stained with DAPI for 5 minutes. Using the Harmony software, the number of nuclei in each well was quantified and used as a proxy for cell survival.

### Patient-derived organoids immunofluorescence

Colonic cancer resection was obtained from Hospital del Mar (Barcelona) with informed consent and the study was approved by the ethical committee. The patient was diagnosed with rectum-sigmoid adenocarcinoma. The isolation of tumor epithelium was performed as described in Sato *et al.*^82^

Colon cancer patient-derived organoids at 10 days of culture were collected and incubated in ice-cold Corning Cell Recovery Solution (Corning; #345235) for 1h at 0°C for Matrigel to dissolve. Organoids were then fixed in 4% Paraformaldehyde (Sigma; #P6148) for 1h at 4°C. After fixation, cells were washed twice with PBS and permeabilized with 1% Triton X-100 for 30 min at room temperature. Cells were blocked 1h at 4°C in PBS + 0.1% BSA (Sigma; #A2058), 0.2% Triton X-100 + 0.1% Tween20. Cells were stained for 48h at 4°C for primary antibodies and 24h for secondary antibodies. The following primary antibodies were used: MTHFD2 (Abcam; #ab151447; 1:200) and VDAC1/Porin (Santa Cruz Biotechnology; #sc-390996; 1:200). The following secondary antibodies were used: goat-anti rabbit 488 (Thermo Fisher Scientific; #A-11008; 1:1000) and goat-anti mouse 555 (Thermo Fisher Scientific; #A-28180; 1:1000). Nuclei were visualized with DAPI (Biogen Cientifica; #BT-40043; 1:1000) and samples mounted with Vectashield Antifade Mounting Medium (Vector Lab; #H1200). Imaging was performed with the Leica SP8 confocal microscope.

### Pull-down – Mass Spectrometry

#### Sample preparation

For pull-down experiments after chromatome fractionation, Protein G Dynabeads (Thermo Scientific; #10004D) were incubated for 6 hours on a rotating wheel at 4°C with primary antibodies MTHFD2 (Abcam; #ab151447; 5 μg) or negative control IgG (Sigma-Aldrich; #I5006; 5 μg). Then, antibody-bound beads were incubated with 2 mg of cytosolic or chromatome extracts overnight on a rotating wheel at 4°C. The complexes were then washed three times with Nuclei Wash Buffer (EDTA 1 mM, IGEPAL 0.1% in PBS). Beads used in the immunoprecipitations were washed three times with 200 mM Ammonium Bicarbonate (ABC) and resuspended in Urea 6M-ABC. Samples were then reduced with dithiothreitol in ABC (30 nM, 37°C, 60 min), alkylated in the dark with iodoacetamide in ABC (60 nM, 25°C, 30 min) and diluted to 1M Urea with 200 mM ABC for trypsin digestion (1 μg, 37°C, overnight shaking, Promega, #V5113). On the next day, beads were separated from the digested extract with a magnet and the peptide mix was acidified with formic acid and desalted with a MicroSpin C18 column (The Nest Group; #SUM SS18V) before LC-MS/MS analysis. Three independent biological replicates for each immunoprecipitation were processed.

#### Chromatographic and mass spectrometric (MS) analysis

Samples were analyzed using an LTQ-Orbitrap Eclipse mass spectrometer (Thermo Fisher Scientific, San Jose, CA, USA) coupled to an EASY-nLC 1200 (Thermo Fisher Scientific (Proxeon), Odense, Denmark). Peptides were loaded directly onto the analytical column and were separated by reversed-phase chromatography using a 50-cm column with an inner diameter of 75 μm, packed with 2 μm C18 particles.

Chromatographic gradients started at 95% buffer A and 5% buffer B with a flow rate of 300 nL/min and gradually increased to 25% buffer B and 75% A in 79 min and then to 40% buffer B and 60% A in 11 min. After each analysis, the column was washed for 10 min with 100% buffer B. Buffer A: 0.1% formic acid in water. Buffer B: 0.1% formic acid in 80% acetonitrile.

The mass spectrometer was operated in positive ionization mode with nanospray voltage set at 2.4 kV and source temperature at 305°C. The acquisition was performed in data-dependent acquisition (DDA) mode and full MS scans with 1 micro scans at a resolution of 120,000 were used over a mass range of m/z 350-1400 with detection in the Orbitrap mass analyzer. Auto gain control (AGC) was set to ‘standard’ and injection time to ‘auto’. In each cycle of data-dependent acquisition analysis, following each survey scan, the most intense ions above a threshold ion count of 10000 were selected for fragmentation. The number of selected precursor ions for fragmentation was determined by the ‘Top Speed’ acquisition algorithm and a dynamic exclusion of 60 sec. Fragment ion spectra were produced via high-energy collision dissociation (HCD) at normalized collision energy of 28% and they were acquired in the ion trap mass analyzer. AGC and injection time were set to ‘Standard’ and ‘Dynamic’, respectively, and isolation window of 1.4 m/z was used.

Digested bovine serum albumin (New England Biolabs; #P8108S) was analyzed between each sample to avoid sample carryover and to assure the stability of the instrument, and QCloud^83^ was used to control instrument longitudinal performance during the project.

#### Data Analysis

Acquired spectra were analyzed using the Proteome Discoverer software suite v1.4 (Thermo Fisher Scientific) and the Mascot search engine^84^ (version 2.6, Matrix Science). The data were searched against a Swiss-Prot human database (as in June 2020) plus a list^85^ of common contaminants and all the corresponding decoy entries. For peptide identification, a precursor ion mass tolerance of 7 ppm was used for the MS1 level, trypsin was chosen as the enzyme and up to three missed cleavages were allowed. The fragment ion mass tolerance was set to 0.5 Da for MS2 spectra. Oxidation of methionine was set as a variable modification whereas carbamidomethylation on cysteine was set as a fixed modification. False discovery rate (FDR) in peptide identification was set to a maximum of 5%. To check the quality of the fractionation, the relative protein abundance of all proteins identified in all the different subcellular compartments was obtained for all the immunoprecipitations. The subcellular location of the proteins was retrieved from Human Protein Atlas^48^ (www.proteinatlas.org). The Significance Analysis of INTeractome (SAINT) express algorithm^49^ was used to score protein-protein interactions. We considered as MTHFD2 potential interactors those with a fold change >= 5 and a Bonferroni False Discovery Rate (BFDR) <= 0.2. The network analysis with MTHFD2 nuclear interactors was performed with Cytoscape^86^ (version 3.9.1) using IntAct^50^ to retrieve protein-protein interactions.

### Pull-down – Western blot

For validation pull-down experiments, cells were washed twice with cold PBS and lysed in Soft-salt Lysis Buffer (10 mM EDTA, 10% glycerol, 0.1% IGEPAL, 50 mM Tris-HCl pH 8 in H_2_O) for 10 minutes on ice. After centrifugation at 3000 rpm for 15 min at 4°C, the supernatant was harvested as the cytosolic fraction, and the nuclear pellet was lysed in High-salt Lysis Buffer (20 mM HEPES pH 7.5, 10% glycerol, 1 mM MgCl_2_, 350 mM NaCl, 0.5% Triton X-100 in H_2_O) for 10 minutes on ice. Nuclear lysates were centrifuged at 13000 rpm at 4°C for 10 min. Balance Buffer (20 mM HEPES pH 7.5, 10 mM KCl, 1 mM MgCl_2_ in H_2_O) was then added to the resulting nuclear supernatant to reach a final NaCl concentration of 150 mM. Nuclear extracts were quantified using Pierce BCA Protein Assay Kit (Thermo Scientific; #PIER23225) and 2 mg of nuclear extract was incubated overnight on a rotating wheel at 4°C with primary antibodies MTHFD2-rabbit (Abcam; #ab151447; 4 μg), MTHFD2-mouse (Abcam; #ab56772; 4 μg), and the negative controls IgG-rabbit (Sigma-Aldrich; #I5006; 4 μg) or IgG-mouse (Sigma-Aldrich; #I5381; 4 μg). Before the addition of the antibodies, 10% of the sample was reserved as input. The next day, the samples were incubated with Protein G Dynabeads (Thermo Scientific; #10004D) for 2 hours at 4°C. The complexes were then washed three times with Wash Buffer (20 mM HEPES pH 7.5, 10% glycerol, 1 mM MgCl_2_, 150 mM NaCl, 0.5% Triton X-100 in H_2_O) and eluted with 2X Laemmli buffer after boiling at 95°C for 5 min.

### Lentiviral production and transfection

HEK293T cells at 70% confluency were used to produce lentiviral particles. Cells were transfected with polyethyleneimine (PEI) (Polysciences; #23966-1) with pCMV-dR8_91 and pVSV-G packaging plasmids, along with the vector of interest, in OptiMeM (Gibco; #11058021). The mixtures with the vectors and PEI were incubated for 5 minutes separately and then mixed and incubated together for 20 minutes to allow complex formation. HEK293T media was changed for serum-free media and the transfection mixture was added dropwise on top. 6 hours after transfection, transfection media was replaced with fresh media. After 48 hours and 72 hours, the viral supernatant was collected and filtered with a 0.45 μm filter unit (Merck Millipore; #051338) and viral aliquots were stored at -80°C until use.

### FUCCI cells generation

To generate stable U2OS and MCF7 cell lines with a Fluorescent Ubiquitination-based Cell Cycle Indicator (FUCCI) system, U2OS and MCF7 cells were transduced with viral particles containing the vectors pLL3.7m-mTurquoise2-SLBP(18-126)-Neomycinin and pLL3.7m-Clover-Geminin(1-110)-IRES-mKO2-Cdt(30-120)-Hygromycin in the presence of polybrene (Sigma-Aldrich; #TR1003G; 10 μg/mL). Since these vectors contained neomycin- and hygromycin-resistance cassettes, respectively, 24 hours after transduction, the media was replaced with fresh media containing 200 μg/mL Geneticin (Thermo Scientific; #10131035) or 150 μg/mL hygromycin (Sigma-Aldrich; #H3274), respectively. Antibiotic selection lasted 4-7 days. Transduced cells were further selected through FACS sorting (BD Influx) to keep cells that showed proper activation and degradation of the FUCCI system. The FUCCI system used is an adaptation of FUCCI4^59^ to show 3 cell cycle-regulated fusion proteins: Clover-Geminin, SLBP-Turquoise2 and Cdt1-mKO2. FUCCI cells were used for immunofluorescence as previously described. The top 5% with the highest MTHFD2 staining in the nucleus or the cytosol were selected for representation.

### MTHFD2 KO and 3xNLS-MTHFD2 knock-in generation

HCT116 cells were nucleofected using the Lonza Amaxa Kit V (Lonza; #VCA-1003) and Amaxa Nucleofector (Lonza) following the HCT116 protocol. Briefly, 2^10^6^ trypsinized cells were resuspended in supplemented nucleofector solution and nucleofected using the D-32 program.

For the MTHFD2 KO, cells were nucleofected with 12 μg of the sgMTHFD2ex4_GW223 cutting vector, which contains Cas9. Two days after nucleofection, single cells were seeded in 96-well plates for isolating single clones. Single clones were tested by Western blot. From the clones tested, we kept two clones that were homozygous KO. These two KO clones were further validated by immunofluorescence and by Sanger sequencing (Eurofins, primers 16-17).

For the 3xNLS-MTHFD2 knock-in, cells were nucleofected with 6 μg of the sgMTHFD2int1-2_GW223 cutting vector (containing Cas9) and 6 μg of the 3xNLS-dTAG-GFP-NEO_GW209 repair vector, following the intron-tagging strategy described by Serebrenik *et al*^87^. Since the repair vector contained a neomycin-resistance cassette, 3 days post-nucleofection cells were treated with 800 μg/mL Geneticin (Thermo Scientific; #10131035) for 7 days. After antibiotic selection, single cells were seeded in 96-well plates for isolating single clones. Single clones were tested by immunofluorescence, and one clone with a homozygous knock-in was kept after Sanger validation (primers 18-19).

### Cellular assays: growth rate, invasion and clonogenic assays

For the growth rate assay, HCT116 MTHFD2 WT, KO and NLS cells were seeded in 12-well plates, fixed with formalin (Sigma-Aldrich; #HT501128) at days 0, 2, 4 and 6, and stained with 0.1% crystal violet solution (Sigma-Aldrich; #HT90132). Wells were then solubilized with 10% acetic acid and measured at 590 nm in a TECAN Infinite M200 Plate Reader.

For the invasion assay, Corning 24-well plates with transwells were used (Corning; #3422). Matrigel (Corning; #354230; 0.3 mg/mL) was seeded in the transwells and left to solidify. After gel formation, HCT116 MTHFD2 WT and KO cells were seeded on the top of the matrigel in serum-free media. After 48 hours, transwells were fixed in 4% formaldehyde (Thermo Fisher Scientific; #28908), permeabilized in 0.1% Triton X-100, 2% BSA in PBS for 15 minutes and stained with DAPI (Sigma-Aldrich; #MBD0015; 1:1000) for 5 minutes. The bottom of the transwells was imaged with a Leica DMI6000B microscope and DAPI mean intensity was used to quantify the number of invading cells.

For the clonogenic assay, HCT116 MTHFD2 WT and KO cells were seeded at a low dilution in 24-well plates. After 15 days, colonies were fixed with formalin (Sigma-Aldrich; #HT501128) and stained with 0.1% crystal violet solution (Sigma-Aldrich; #HT90132). The plates were scanned, and the area covered by colonies was obtained with the ImageJ plugin ColonyArea^88^.

### RNA-sequencing

#### Sample preparation

Three biological replicates were obtained from 2.5^10^6^ HCT116 MTHFD2 WT and KO1 cells at an early time-point and after two months of cell culture. RNA was extracted using the PureLink RNA mini kit (Thermo Fisher Scientific; #12183018A). Libraries were prepared from 500 ng of RNA using the TruSeq stranded mRNA Library Prep (Illumina; #20020594) according to the manufacturer’s protocol, to convert total RNA into a library of template molecules of known strand origin and suitable for subsequent cluster generation and DNA sequencing. Final libraries were analyzed using the Bioanalyzer DNA 1000 (Agilent; #5067-1504) to estimate the quantity and validate the size distribution and were then quantified by qPCR using the KAPA Library Quantification Kit KK4835 (Roche; #07960204001) before the amplification with Illumina’s cBot. Libraries were sequenced on the Illumina HiSeq 2500 machine using single-read 50bp sequencing.

#### Data Analysis

Quality control was performed with FastQC^89^ (version 0.11.9). Single-end, 50-bp-long reads were aligned to the GRCh38.p13 Homo Sapiens reference genome using the STAR Aligner^90^ (version 2.7.6a). Gene level counts were obtained with featureCounts from subread^91^ (version 2.0.1), using gene annotations downloaded from Gencode (Release 38 GRCh38.p13). Differential expression analysis was performed in R (version 4.1.1) using the DESeq2 package^92^ (version 1.32.0). Principal Component analysis was performed with the function prcomp. The lfcShrink function from DESeq2 with the apeglm method^93^ was used for visualization purposes. Genes with adjusted p-value < 0.05 were considered differentially expressed. Those differentially expressed genes that were shared in both time points, and with an absolute log_2_ Fold Change > 0.58 in at least one time point were further considered for functional enrichment analysis. Gene Ontology enrichment analysis was performed using the ClusterProfiler^94^ package (version 4.0.5). Gene Set Enrichment Analysis (GSEA) was performed with GSEA software^95^ (version 4.1.0) using the ‘hallmark’ gene set from the MSigDB collection. For the centromeric analysis, centromeric coordinates of the assembly GRCh38 were retrieved from the Table Browser of UCSC (University of California Santa Cruz) and the intersect was obtained with bedtools^96^.

### RNA Extraction and RT-qPCR

RNA was extracted using the PureLink RNA Mini Kit (Thermo Fisher Scientific; #12183018A) and converted into cDNA using the High-Capacity RNA-to-DNA kit (Applied Biosystems; #4387406). Quantitative PCR was performed using the Power SYBR Green PCR Master Mix (Applied Biosystems; #4367659) in a ViiA 7 Real-Time PCR System (Thermo Fisher Scientific). Results were analyzed with the Design and Analysis Software QuantStudio 6/7 Pro systems (Thermo Fisher Scientific, version 2.6). Primers 20-59 (Supplementary Methods Table 2) were used, obtained from Contreras-Galindo *et al.*^97^

### Chromatin Immunoprecipitation (ChIP)-qPCR

20^10^6^ HCT116 WT and KO1 cells were crosslinked by adding on the plate culture media with 1% formaldehyde (Thermo Fisher Scientific; #28908) for 10 min at room temperature shaking. Crosslinking was stopped by adding glycine at a final concentration of 0.125 M for 5 min at room temperature shaking. Crosslinked cells were washed twice with PBS, scrapped in PBS with proteinase inhibitors (Roche; #4693132001) and collected by centrifugation for 5 min at 4000 rpm 4°C. Cell pellets were lysed in Nuclei Lysis Buffer (10 mM EDTA, 1% SDS, 50 mM Tris-HCl pH 8 in H_2_O) supplemented with proteinase inhibitors. Nuclear extracts were sonicated on a Bioruptor Pico (Diagenode) for 10 cycles 30 sec ON/30 sec OFF in 1.5 mL Diagenode tubes (Diagenode; #C30010016) to generate 100–400 bp DNA fragments. Sonicated extracts were centrifuged for 15 minutes at 13000 rpm 4°C to remove insoluble material, and the supernatant was kept and diluted 1:10 in Immunoprecipitation Buffer (1.2 mM EDTA, 167 mM NaCl, 16.7 mM Tris-HCl pH 8, 1.1% Triton X-100 in H_2_O) supplemented with proteinase inhibitors. 20 μg of chromatin were used for each immunoprecipitation, which were incubated overnight on a rotatory wheel at 4°C with the primary antibodies H4K20me1 (Diagenode; #C15410034; 2 μg), H3K9me3 (Diagenode; #C15410193; 2 μg), H3K27me3 (Diagenode; #C15410195; 2 μg) and IgG-rabbit (Sigma-Aldrich; #I5006; 2 μg) as a negative control. Before the addition of the antibodies, 10% of each sample was reserved as input. The next day, the samples were incubated with Protein A Dynabeads (Thermo Scientific; #10002D) for 2 hours at 4°C. The complexes were then washed four times with four wash buffers: Wash Buffer 1 (2 mM EDTA, 150 mM NaCl, 0.1% SDS, 20 mM Tris-HCl pH 8, 1% Triton X-100 in H_2_O), Wash Buffer 2 (2 mM EDTA, 400 mM NaCl, 0.1% SDS, 20 mM Tris-HCl pH 8, 1% Triton X-100 in H_2_O), Wash Buffer 3 (1 mM EDTA, 1% IGEPAL, 0.25 M LiCl, 1% NaDOC, 10 mM Tris-HCl in H_2_O) and TE Buffer (1 mM EDTA, 10 mM Tris-HCl in H_2_O). Beads were resuspended in 200 μL of freshly-prepared ChIP Elution Buffer (0.1 M NaHCO_3_, 1% SDS in H_2_O) and incubated at 65°C for 1 hour. Then, beads were removed from the chipped chromatin with the magnet, and the immunoprecipitated chromatin was de-crosslinked by adding 200 mM NaCl and incubating at 65°C overnight. On the next day, RNA and protein were digested in 40 mM Tris-HCl pH 6 and 10 mM EDTA by adding first RNAse A (Qiagen; #19101, 1.5 hours, 37°C), followed by Proteinase K (Thermo Fisher Scientific; #EO0491, 1 hour, 45°C). Inputs were incubated in parallel during the de-crosslinking step. Finally, DNA was purified using a MiniElute PCR Purification Kit (Qiagen; #28006) and eluted in nuclease-free water, and quantitative PCR was performed using the Power SYBR Green PCR Master Mix (Applied Biosystems; #4367659) in a ViiA 7 Real-Time PCR System (Thermo Fisher Scientific). Results were analyzed with the Design and Analysis Software QuantStudio 6/7 Pro systems (Thermo Fisher Scientific, version 2.6). Primers 20-23,36-39 (Supplementary Methods Table 2) were used, obtained from Contreras-Galindo *et al.*^97^

### Nanopore whole-genome sequencing

#### Genomic DNA extraction

3x10^^6^ HCT116 WT and KO1 frozen cell pellets were used to extract high molecular weight genomic DNA (HMW gDNA) following the manufacturer’s protocol of Nanobind tissue kit (Circulomics). The HMW gDNA eluate was quantified by Qubit DNA BR Assay kit (Thermo Fisher Scientific) and the DNA purity was evaluated using Nanodrop 2000 (Thermo Fisher Scientific) UV/Vis measurements. The HMW gDNA samples were stored at 4°C.

#### Long Read Whole-genome library preparation and sequencing

After quality control of the HMW gDNA for purity, quantity and integrity for long-read sequencing, the libraries were prepared using the 1D Sequencing kit SQK-LSK110 from Oxford Nanopore Technologies (ONT). Briefly, 4.0 μg of the DNA were DNA-repaired and DNA-end-repaired using the NEBNext FFPE DNA Repair Mix (New England Biolabs; #M6630) and the NEBNext UltraII End Repair/dA-Tailing Module (New England Biolabs; #E7546), and followed by the sequencing adaptors ligation, purified by 0.4X AMPure XP beads (Agencourt, Beckman Coulter) and eluted in Elution Buffer. The sequencing runs were performed on PromethIon 24 (ONT) using a flow cell R9.4.1 FLO-PRO 002 (ONT) and the sequencing data was collected for 110 hours. The quality parameters of the sequencing runs were monitored by the MinKNOW platform (version 21.11.7) in real-time and basecalled with modified basecalling for 5mC using Guppy (version 5.1.13).

#### Basecalling and mapping

Raw Nanopore data in the format of fast5 files were analyzed using Master of Pores 2^98^ suite performing the basecalling with Guppy (version 6.1.1) and the modified base model “dna_r9.4.1_450bps_modbases_5mc_cg_hac_prom”. The unaligned bam outputs were first converted to fastq files keeping the information about the modified bases in the header, filtered using nanoq^99^ for removing the reads with average coverage lower than 7, and then aligned to the human T2T (telomere to telomere) genome using Winnowmap^100^ (parameters -y -ax map-ont), a minimap-derived aligner specifically designed for repetitive regions. Final alignment files were sorted, indexed and filtered with samtools^101^ for further analysis. Only primary alignments were retained.

#### Methylation analysis

From the final filtered alignment bam files, methylation data (counts of reads of methylated and unmethylated CpG) were obtained with modbam2bed (parameters -extended, -aggregate, -cpg) in bedmethyl format. R package methylKit^102^ (version 1.20.0) and custom scripts were used to analyze methylation data and transform between different formats. The significance of methylation level change between the conditions, for every CpG site, was calculated with methylKit. Those sites with the largest methylation variation were extracted, by filtering on methylation difference >= 15% and q-value < 0.05. Centromeric coordinates for each chromosome were acquired from Altemose *et al.*^103^

#### Structural variation analysis

Aligned reads were fed to CuteSV tool^104^ for predicting large structural variations. These predicted variants were annotated using Ensembl Variant Effect Predictor (VEP)^105^. Variants were crossed with methylation information using bedtools^96^. Plots were made using scripts written with R statistical language (version 4.1.1).

### Chromosome Banding Analysis of cell lines

Cytogenetic studies were performed in HTC116 WT and KO cells. Cell cultures were incubated for 3-4 days at 37 °C/5% CO_2_ up to the time of its extraction, when the cell culture was 80% confluent. 10 μg/mL of colcemid were added to the cultures and incubated for 4 hours for arresting the cells in metaphase. After trypsinization, cells were swollen hypotonically and fixed with Carnoy (methanol:acetic acid). Then, the cells were dropped on 4-5 slides. For chromosome banding, slides with metaphase spreads were treated in a slide warmer at 100°C for 1 hour and stained with Wright’s solution to create characteristic light and dark bands. Metaphase spreads were captured using an automated imaging system for cytogenetics (CytoInsight GSL, Leica Biosystems) and karyotyped with Cytovision Software 7.0 (Applied Imaging Corporation). Karyotypes were described following the International System for Human and Cytogenetic Nomenclature (2020). A minimum of 20 metaphases were analyzed.

### Cell cycle analysis

HCT116 MTHFD2 WT and KO cells were seeded on 6-well plates and treated the day after either with dimethyl sulfoxide (DMSO) (PanReac AppliChem; #A3672) or CDK1 inhibitor RO-3306 9μM (MedChem Express; #HY-12529) during 20 hours to synchronize cells at the end of G2 phase. Treated cells were then washed three times with PBS and released in normal media. For cells treated with DMSO, cells were harvested immediately after release. For cells treated with RO-3306, cells were harvested at 0, 0.5, 1, 1.5, 2 and 2.5 hours after release. Cells were then fixed with 1 mL of 70% cold ethanol in PBS, added dropwise while vortexing, left on ice for 2 hours and kept overnight at 4 °C. Cells were then washed four times with 5 mM EDTA in PBS, and stained with propidium iodide solution (15 μg/mL propidium iodide (Life Technologies; #P3566), 1 mM sodium citrate, 30 μg/mL RNAse A (Qiagen; #19101)) overnight at 4°C in the dark. Cells were analyzed in the flow cytometry analyzer LSRII (BD Biosciences) and plotted with FlowJo (version 10.8.2).

### Whole-genome CRISPR genetic screening

#### Sample preparation

The human CRISPR knockout pooled (Brunello) library^106^ consists of 76,441 sgRNAs targeting 19,114 genes (whole genome), with ∼4 sgRNA/gene, as well as 1000 intergenic control sgRNAs in a Cas9-expressing lentiviral vector. The Brunello library was amplified with the QIAGEN Plasmid Plus Mega Kit (Qiagen; #12981). For virus production, HEK293T cells were transfected with the Brunello lentiCRISPRv2 plasmid as previously described. HCT116 WT and KO cells were infected with the lentiviral particles containing the Brunello library at a multiplicity of infection (MOI) of 0.4 in the presence of polybrene (Sigma-Aldrich; #TR1003G; 10 μg/mL). The next day, puromycin-containing medium (Sigma-Aldrich; #P8833; 2 μg/mL) was added to select transductants. At 8 days post-selection, half of the cells were harvested as the initial population and the other half were reseeded, maintaining a 500X coverage. Cells were kept in culture for three weeks, when the final population was harvested. Genomic DNA from all samples was extracted using the QIAmp DNA Blood Midi kit (Qiagen; #51106) and treated with RNAse A (Qiagen; #19101). The sgRNA library was prepared by a first PCR with the Phusion High Fidelity DNA Polymerase (Thermo Fisher Scientific; #F530) with a mixture of P5 forward primers with staggers from 3 to 6 bp and a P7 reverse primer (Primers 60-64, Supplementary Methods Table 2). The number of cycles was optimized for each sample to prevent over-amplification, and the DNA input for each sample corresponded to a coverage of ∼500X. A second PCR was performed using NEBNext Q5 Hot Start HiFi PCR Master Mix (New England Biolabs; #M0543) and Index 1 (i7) and Index 2 (i5) to complete the adaptors and to add the barcodes. Final libraries were analyzed using Agilent Bioanalyzer (Agilent; #5067-4626) to estimate the quantity and check size distribution, and were then quantified by qPCR using the KAPA Library Quantification Kit (KapaBiosystems; #KK4835) prior to amplification with Illumina’s cBot. Samples were sequenced on the Illumina HiSeq 2500 machine using single-read 50bp sequencing.

#### Data Analysis

MAGeCK was used for alignment, gRNA count, copy number variation (CNV) correction, and for the obtention of gene-level depletion and enrichment scores^67^. MAGeCKFlute^107^ was additionally used to correct for cell cycle-related effects between MTHFD2 WT and KO cells. Gene log_2_ fold change was calculated by taking the average of the log_2_ fold change for all sgRNAs targeting the same gene.

### Etoposide treatment

HCT116 WT and KO cells were seeded in 96-well plates and, after 24 hours, cells were treated with either DMSO (PanReac AppliChem; #A3672) or different doses of etoposide (MedChem Express; #HY-13629): 0.25, 0.5, 1, 2 and 4 μM for 72 hours. Afterward, cells were fixed with formalin (Sigma-Aldrich; #HT501128) and stained with 0.1% crystal violet solution (Sigma-Aldrich; #HT90132). Cells were then solubilized with 10% acetic acid and measured at 590 nm in a TECAN Infinite M200 Plate Reader.

### Folate metabolites supplementation

HCT116 WT and KO cells were seeded in 96-well plates and, after 24 hours, cells were supplemented with either DMSO (PanReac AppliChem; #A3672) or different metabolites: formic acid (low dose 0.25 mM, high dose 1 mM, Thermo Fisher Scientific; #A11750), folate (low dose 0.125 μM, high dose 50 μM, MedChem Express; #HY-1663), 5,10-methylenetetrahydrofolate (low dose 0.25 μM, high dose 2.5 μM, MedChem Express; #HY-14769) or SAM (low dose 0.375 μM, high dose 50 μM, MedChem Express; #HY-B0617A) for 72 hours. Afterward, cells were fixed for immunofluorescence to measure the mitotic index as previously described.

### TH9619 treatment

HCT116 WT cells were seeded in 96-well plates and, after 24 hours, cells were treated with DMSO (PanReac AppliChem; #A3672) or 25, 62.5 and 156.25 nM TH9619 inhibitor (One-Carbon Therapeutics) for 96 hours. After TH9619 inhibitor treatment, cells were fixed for immunofluorescence to measure the mitotic index and H4K20me1 levels. To measure the sensitivity of HCT116 cells to TH9619, cells were seeded in 96-well plates and, after 24 hours, cells were treated with DMSO (PanReac AppliChem; #A3672) or 4 nM, 13 nM, 41 nM, 123 nM, 370 nM, 1.11 μM, 3.33 μM, 10 μM and 30 μM of TH9619 inhibitor for 96 hours. After TH9619 inhibitor treatment, cells were fixed for immunofluorescence to quantify DAPI staining.

### TCGA RNA-sequencing data analysis

RNA-sequencing data from 31 tumor types belonging to 26 different primary sites from The Cancer Genome Atlas (TCGA) database were retrieved using the Genomic Data Commons (GDC) data portal from The National Cancer Institute^46^. For assessing MTHFD2 expression in tumor versus healthy tissues, FPKM (fragments per kilobase million) values of the *MTHFD2* gene were first converted to TPM (transcripts per million) values. 15 solid tumor types, where paired normal tissue data was available and with a minimum of 10 samples, were kept. In these tumor types, tumor and healthy MTHFD2 expression were compared with a paired two-tailed Wilcoxon test in R (version 4.1.1).

For the machine learning analysis, tumor and healthy MTHFD2 expression from breast (BRCA), lung (LUAD and LUSC) and colon (COAD) cancer, separately, were used to predict the sample status: healthy or tumor. Two-thirds of the data were used for training a random forest machine learning algorithm with a number of trees of 500 using the randomForest package^108^ (version 4.7-1.1) in R. The resting one-third of the data was used to evaluate model performance and to obtain the ROC (receiver operating characteristic) curves and the corresponding AUC (area under the curve) values, using the packages caret^109^ (version 6.0-93) and ROCR^110^ (version 1.0-11).

For the co-expression analysis, for each cancer type, Pearson correlation was measured between the TPM expression values of each one of the enzymes of the folate metabolism (ATIC, DHFR, GART, MTHFD1, MTHFD1L, MTHFD2, MTHFD2L, MTHFR, MTR, SHMT1, SHMT2) and all the other genes. *P*-values were adjusted with the Benjamini & Hochberg correction. Positively correlated genes with a Pearson correlation coefficient (r) > 0.6 and *p*-adjusted values < 0.05 were further considered, and those which were positively correlated with the folate enzyme in >= 10 cancer types were selected for over-representation analysis. Gene Ontology enrichment analysis was performed using the ClusterProfiler^94^ package (version 4.0.5) in R.

### CCLE data analysis

MTHFD2 RNA levels (from Expression Public 22Q4), MTHFD2 protein levels (from Proteomics) and MTHFD2 essentiality levels (from CRISPR DepMap Public 22Q4+Score) were retrieved from the Cancer Cell Line Encyclopedia (CCLE) data^47^ hosted on the DepMap Portal (https://depmap.org). The aneuploidy score and aneuploidy group classification from all CCLE cell lines were retrieved from Cohen-Sharir *et al.*^66^

### Proteome-HD data analysis

The ProteomeHD tool^57^, which employs proteomics data in response to biological perturbations to perform co-regulation analysis using unsupervised machine learning, was used to identify proteins that are co-regulated with MTHFD2. The 5% strongest co-regulated proteins were selected, along with their corresponding enriched Gene Ontology Biological Process terms.

### MTHFD2 levels in cell cycle phases

The MTHFD2 SILAC ratios of three biological replicates in each of the cell cycle phases (G1, S, G2, M) were obtained from the supplementary file 2 from Ly *et al.*^58^ As mentioned in their publication, for each biological replicate, the ratios were normalized to the ratio measured in G1, and an offset was then added to the G1 ratio to account for the difference in time between cell division and an average G1 cell. Statistical significance was obtained using a one-sample two-tailed *t*-test.

### Statistical analysis

All the statistical parameters including the exact value of the number of replicates, number of cells, deviations, *p*-values and type of statistical test are reported in their respective figures. Statistical analysis was performed across biological replicates, by taking the average of the respective technical replicates, when appropriate. Statistical significance was analyzed using unpaired two-tailed Student’s *t-*test after testing for normality with Shapiro-Wilk test and equal variance with Levene test. If assumption of normality and homoscedasticity were not fulfilled, the unpaired (or paired, when relevant) non-parametric two-tailed Wilcoxon test was used. In all cases, ns: not significant (*P* > 0.05), *: *P* < 0.05, **: *P* < 0.01, ***: *P* < 0.001, ****: *P* < 0.0001.

### Data availability

The raw proteomics data have been deposited to the PRIDE^111^ repository with the dataset identifier PXD041821.

Sequencing samples (raw data and processed files) are available at NCBI GEO. RNA-sequencing data is under the accession number GSE230750.

Genome-wide CRISPR genetic screening raw data have been deposited to the European Nucleotide Archive (ENA) under the accession number PRJEB61666.

### Code availability

All the scripts used for this manuscript are publicly available in the GitHub repository https://github.com/NataliaPardoL/MTHFD2_mitosis.

## Acknowledgments

We would like to thank the CRG and CNAG Sequencing Facilities (Barcelona, Spain) for all next generation sequencing, the CRG Proteomics Facility (Barcelona, Spain) for the MTHFD2 interactome analysis, the CRG Flow Cytometry Facility (Barcelona, Spain) for the sorting, the IMIM Platform for molecular cytogenetics (Barcelona, Spain) for the karyotype analysis, and MARbiobank for provision of patient samples for organoid generation. The CRG/UPF Proteomics Unit is part of the Spanish Infrastructure for Omics Technologies (ICTS OmicsTech).

NPL is supported by a Boehringer Ingelheim Fonds Ph.D. fellowship. The Sdelci lab’s contributions to this study were funded by an ERC Starting Grant (ERC-StG-852343-EPICAMENTE) and by a Spanish Plan Estatal grant (Ministerio de Ciencia e Innovación, PID2019-110598GA-I00-COMBAT)

## Author Contributions

S.S. and N.P.L. conceptualized and designed the project. N.P.L. performed and/or analyzed all the experiments shown in Fig. 1-5 and Extended Data Fig. 1-10. S.S. performed the mitotic phases and mitotic defects quantification shown in Fig. 4-5 and Extended Data Fig. 7. A.G. and L.C. performed the analysis of Nanopore whole-genome sequencing data shown in Fig. 3 and Extended Data Fig. 6. A.G.Z. performed the FUCCI experiments shown in Fig. 2 and Extended Data Fig. 4. L.E. cloned of all the *de novo* vectors described in this manuscript. L.G.L. contributed to the pull-down and ChIP-qPCR experiments shown in Extended Data Fig. 2 and Fig. 3. R.G.A. performed the qPCR experiments shown in Fig. 3. E.D. contributed to the pull-down experiments shown in Extended Data Fig. 2. J.S. performed the immunofluorescence of patient-derived colon organoids shown in Extended Data Fig. 1. S.S., J.P. and L.B.M. supervised the study. S.S. acquired the funding. S.S. and N.P.L. wrote the manuscript with contributions from all the authors.

---

**Extended Data Fig. 1:**
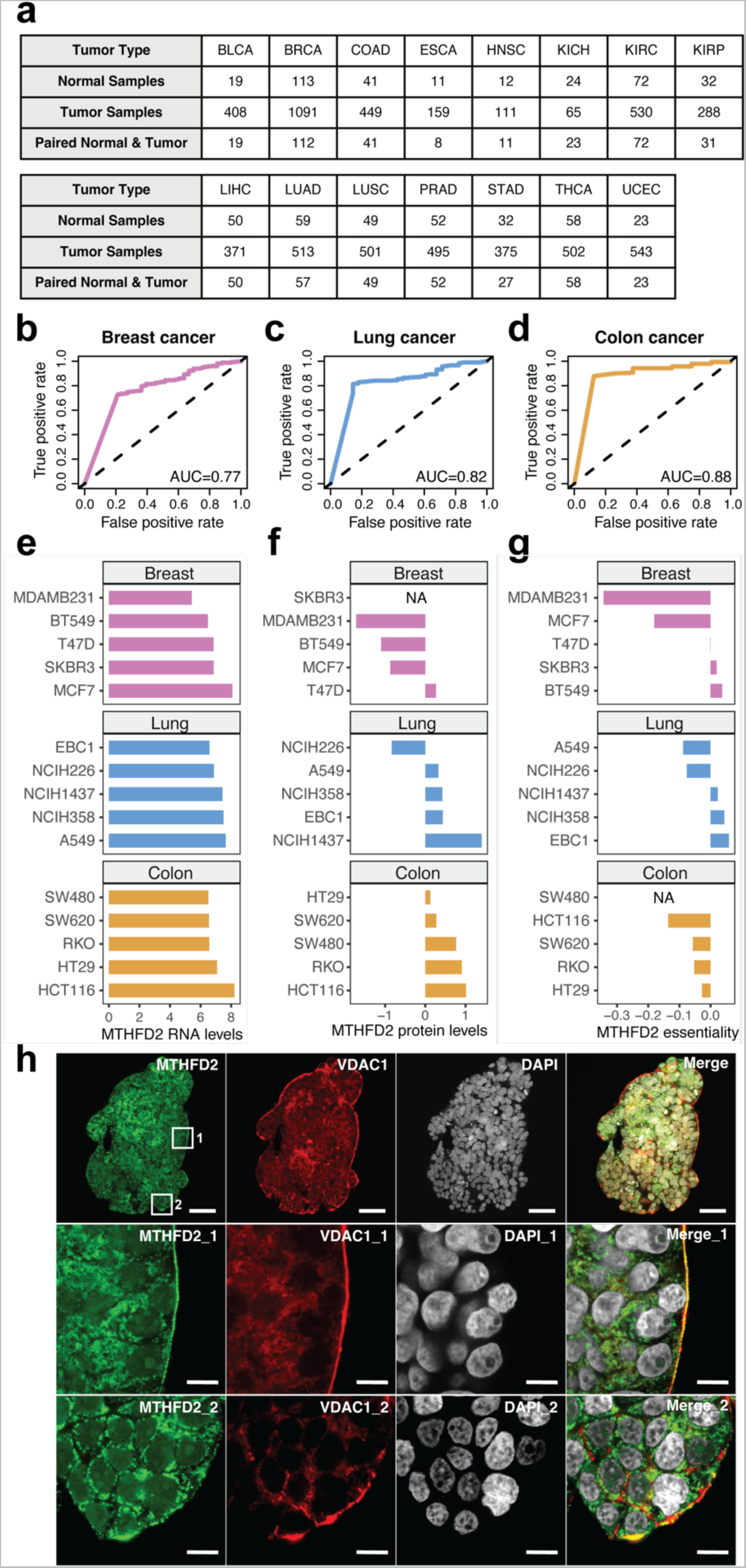
MTHFD2 expression level is a potent cancer predictor. **a**, Schematic table with the number of normal and tumor RNA-sequencing data per tumor type obtained from The Cancer Genome Atlas (TCGA) database. BLDA, bladder urothelial carcinoma; BRCA, breast invasive carcinoma; COAD, colon adenocarcinoma; ESCA, esophageal carcinoma; HNSC, head and neck squamous cell carcinoma; KICH, kidney chromophobe; KIRC, kidney renal clear cell carcinoma; KIRP, kidney renal papillary cell carcinoma; LIHC, liver hepatocellular carcinoma; LUAD, lung adenocarcinoma; LUSC, lung squamous cell carcinoma; PRAD, prostate adenocarcinoma; STAD, stomach adenocarcinoma; THCA, thyroid carcinoma; UCEC, uterine corpus endometrial carcinoma. **b-d,** Random forest model performance in the form of ROC (receiver operating characteristic) curves with the corresponding AUC (area under the curve) values for breast (b), lung (c) and colon (d) cancer. **e-g,** MTHFD2 RNA levels (e), protein levels (f) and essentiality values (g) in a panel of cancer cell lines from breast, lung and colon cancer. NA, data not available. **h,** Immunofluorescence of colon cancer-patient organoids. MTHFD2 is shown in green, VDAC1 (mitochondrial marker) in red and DAPI in grey; confocal mode; scale bar 50 μm for the global image and 10 μm for the zoomed images.

**Extended Data Fig. 2:**
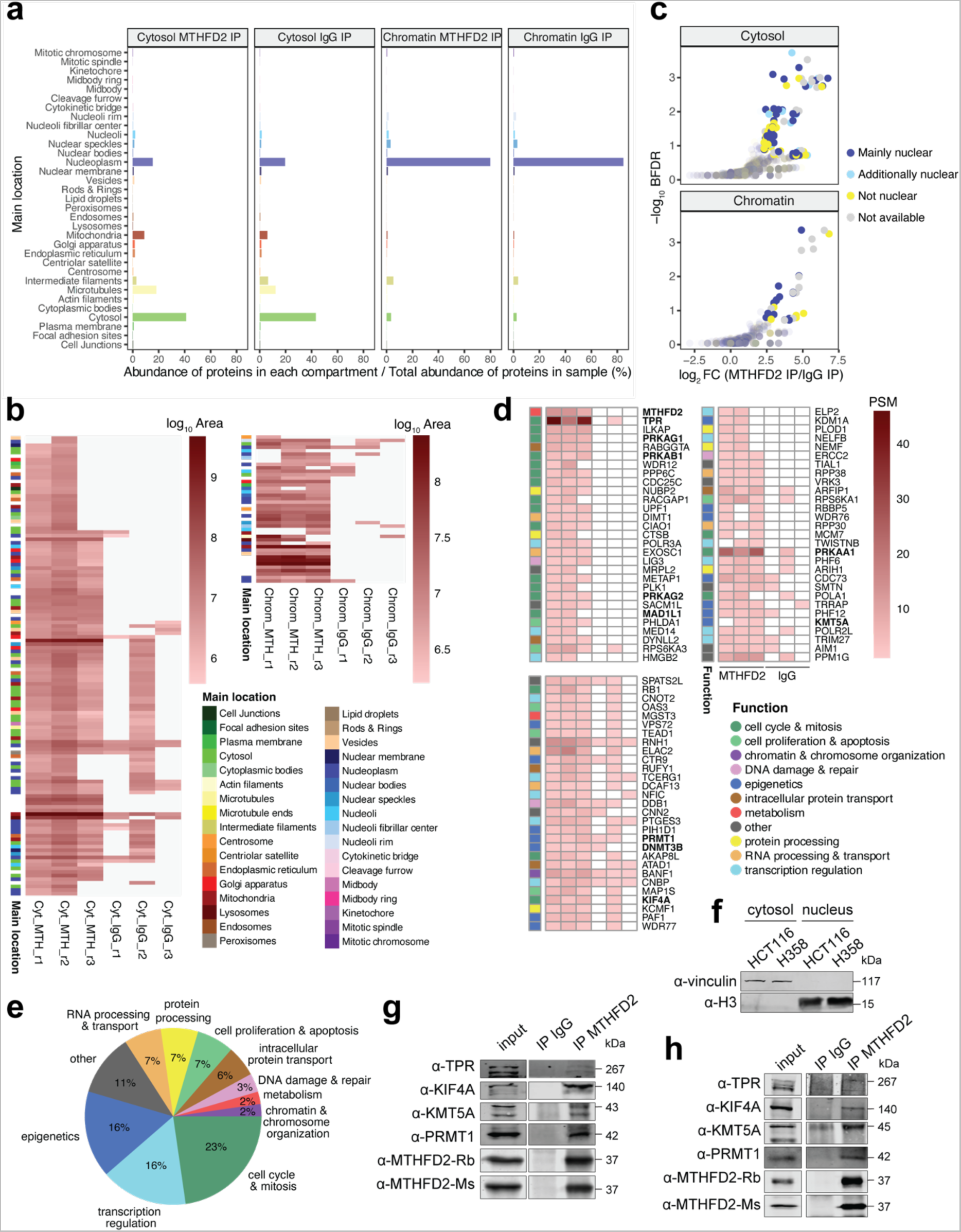
Quality control and validation of the MTHFD2 interactome. **a**, Normalized abundance of detected proteins in each subcellular compartment in the cytosolic MTHFD2, cytosolic IgG, chromatin MTHFD2, chromatin IgG immunoprecipitations (IP) performed in HCT116 cells. **b,** Heatmap of the log_10_ area of the putative cytosolic and chromatin interactors with a log_2_ fold change (FC) >= 2.3 and Bonferroni False Discovery Rate (BFDR) <= 0.2, colored by their subcellular compartment. Cyt, cytosol; Chrom, chromatin; MTH, MTHFD2; r, replicate. **c,** Volcano plots of putative cytosolic (top) and chromatin (bottom) interactors with a log_2_ fold change (FC) >= 2.3 and Bonferroni False Discovery Rate (BFDR) <= 0.2, colored by their nuclear localization. **d,** Heatmap of the peptide spectrum matches (PSM) of the top nuclear MTHFD2 interactors, colored by their functional category. **e,** Pie chart of the functional categories of the top MTHFD2 nuclear interactors. **f,** Western blot of cytosolic and nuclear extracts of HCT116 and H358 cells. Vinculin and histone H3 are used as cytosolic and nuclear markers, respectively. **g-h,** Western blot of nuclear input fractions and immunoprecipitations (IP) with IgG or MTHFD2 in HCT116 (g) and H358 (h) cells.

**Extended Data Fig. 3:**
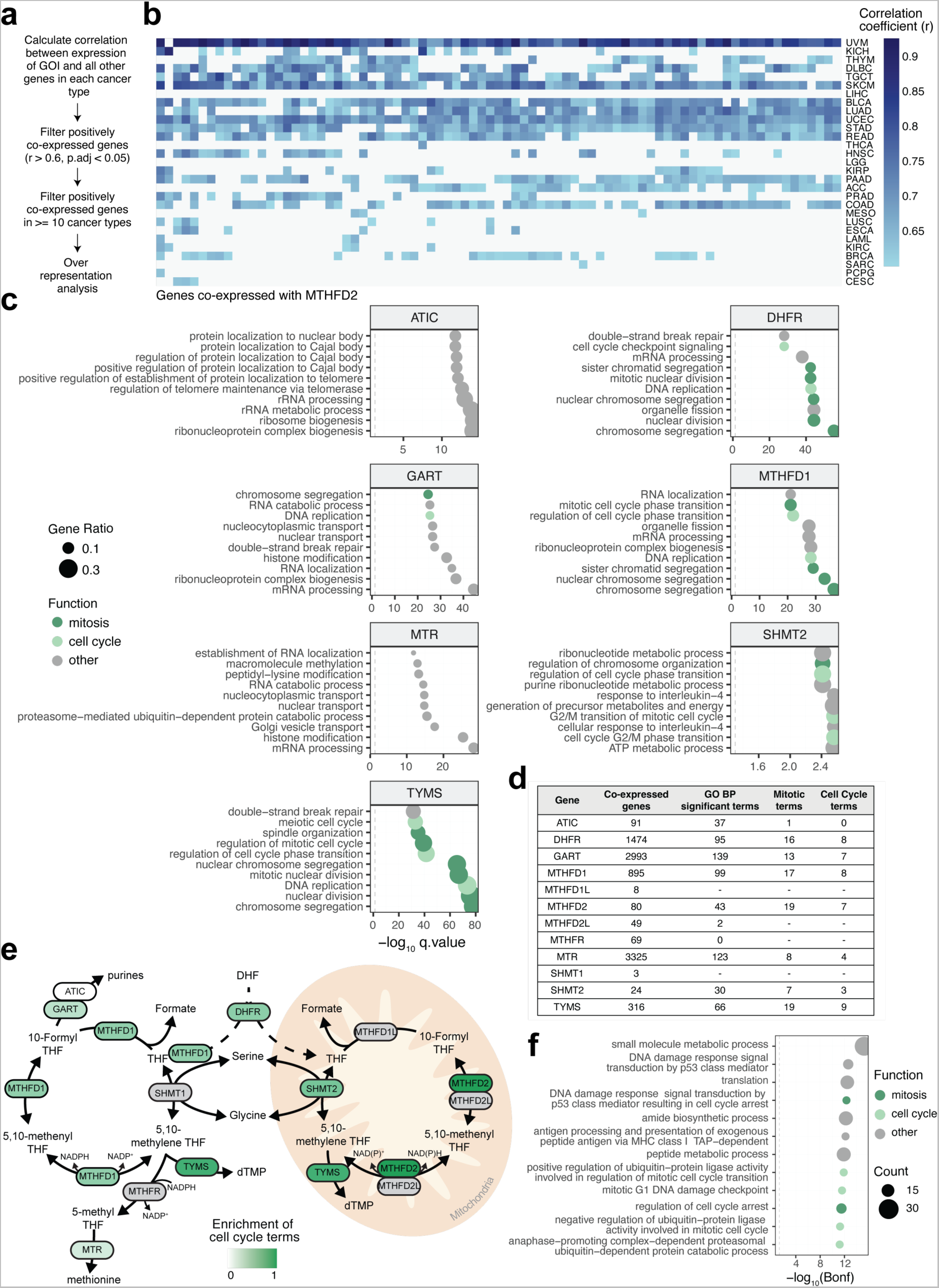
MTHFD2 is co-expressed with mitotic genes. **a**, Schematic of the co-expression analysis. GOI, gene of interest; p.adj, p-value adjusted. **b,** Heatmap of the correlation coefficient (r) of genes co-expressed with MTHFD2 (r > 0.6 and p-value adjusted < 0.05) it at least 10 different cancer types. ACC, adrenocortical carcinoma; BLDA, bladder urothelial carcinoma; BRCA, breast invasive carcinoma; CESC, cervical squamous cell carcinoma and endocervical adenocarcinoma; COAD, colon adenocarcinoma; DLBC, diffuse large B-cell lymphoma; ESCA, esophageal carcinoma; HNSC, head and neck squamous cell carcinoma; KICH, kidney chromophobe; KIRC, kidney renal clear cell carcinoma; KIRP, kidney renal papillary cell carcinoma; LAML, acute myeloid leukemia; LGG, lower grade glioma; LIHC, liver hepatocellular carcinoma; LUAD, lung adenocarcinoma; LUSC, lung squamous cell carcinoma; MESO, mesothelioma; PAAD, pancreatic adenocarcinoma; PCPG, pheochromocytoma and paraganglioma; PRAD, prostate adenocarcinoma; READ, rectum adenocarcinoma; SARC, sarcoma; SKCM, skin cutaneous melanoma; STAD, stomach adenocarcinoma; TGCT, testicular germ cell tumors; THCA, thyroid carcinoma; THYM, thymoma; UCEC, uterine corpus endometrial carcinoma; UVM, uveal melanoma. **c,** Top terms of Biological Process Gene Ontology enrichment of folate enzymes co-expressed genes. The dashed line indicates the threshold of q.value = 0.05. **d,** Schematic table showing the number of co-expressed genes obtained per each of the one-carbon folate metabolism enzymes, the number of Gene Ontology (GO) biological process (BP) significant terms obtained from the co-expressed genes, and from this, the number of terms which were related to mitosis or cell cycle. **e,** One-carbon folate metabolism pathway with enzymes colored by the enrichment of cell cycle terms in the Gene Ontology enrichment analysis performed with their core co-expressed genes. Gray indicates the absence of data. DHF, dihydrofolate; THF, tetrahydrofolate. Scheme adapted from Lin *et al*^4^. **f,** Gene Ontology Biological Process terms enriched in co-regulated proteins with MTHFD2. Bonf, Bonferroni False Discovery Rate. The dashed line indicates the threshold of Bonf = 0.05.

**Extended Data Fig. 4:**
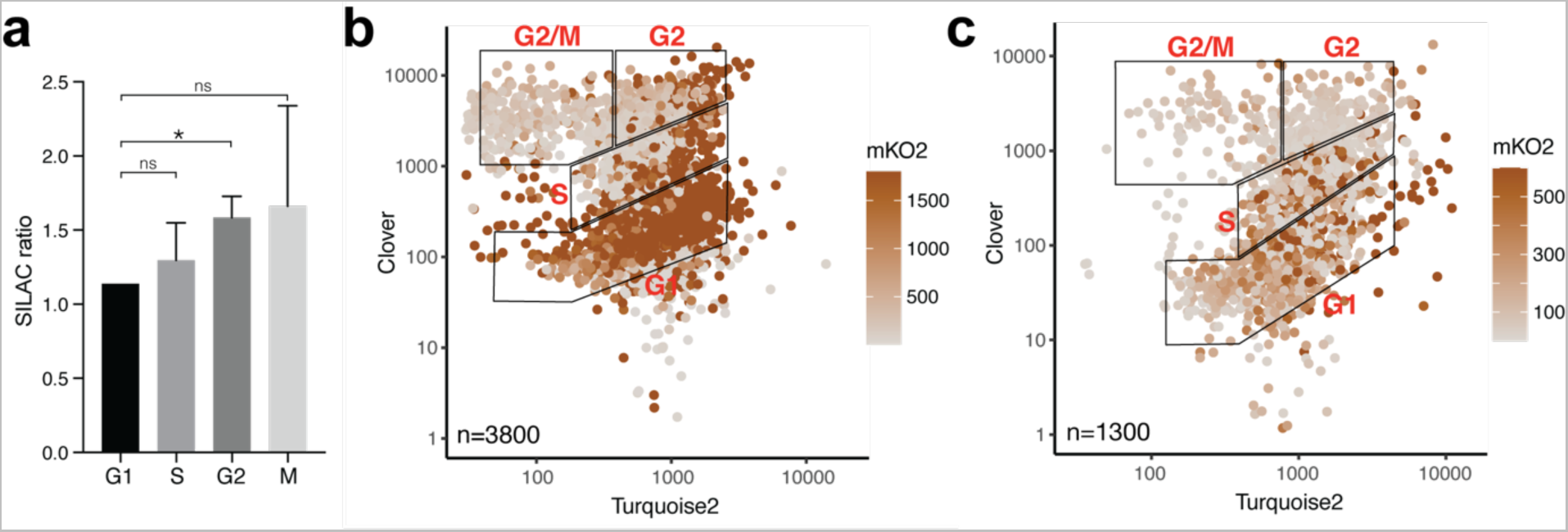
MTHFD2 levels differ across cell cycle phases. **a**, SILAC ratio of MTHFD2 in G1, S, G2 and M phases in NB4 cells from the data obtained in Ly *et al.*^58^; means + s.d. (*n*=3), one-sample two-tailed *t*-test (ns, non-significant; *, *P*<0.05). **b-c,** FUCCI-adapted MCF7 (b) and U2OS (c) cells along with the cell cycle phase; turquoise and clover mean intensities (x and y axis, respectively) are in log_10_ scale.

**Extended Data Fig. 5:**
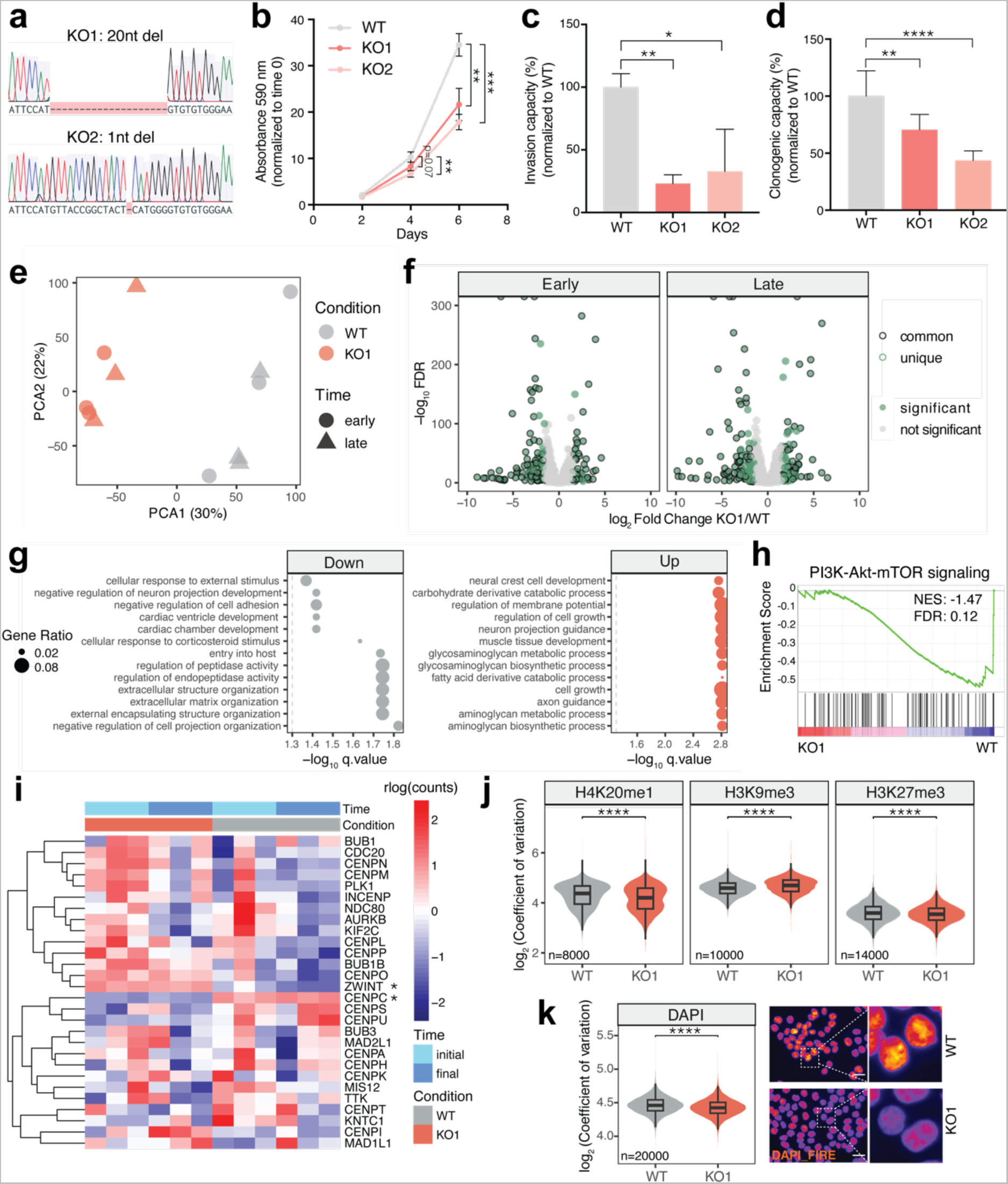
The function of MTHFD2 in mitosis is not transcriptionally mediated. **a**, Sequence validation of HCT116 MTHFD2 knock-out (KO1 and KO2) cells by Sanger sequencing. **b,** Growth curve of MTHFD2 wild-type (WT), KO1 and KO2 cells normalized to time 0 days measured as crystal violet absorbance at 590 nm; means ± s.d. (*n*=3), unpaired two-tailed *t*-test (**, *P*<0.01; ***, *P*<0.001). **c-d,** Invasion (c) and clonogenic (d) capacity of WT, KO1 and KO2 cells normalized to the WT condition; means + s.d. (*n*=3), unpaired two-tailed *t*-test (*, *P*<0.05; **, *P*<0.01; ****, *P*<0.0001). **e,** Principal Component Analysis (PCA) of transcriptomics data from WT and KO1 cells at early (right after KO generation) and late (after two months in cell culture) time points. **f,** Volcano plot of differentially expressed genes between KO1 and WT conditions at both time points. Genes with absolute log_2_ fold change > 1.5 and False Discovery Rate (FDR) < 0.05 are shown. Shared genes between both MTHFD2 KOs are indicated with a black stroke. **g,** Gene Ontology Biological Process enrichment analysis performed with shared down- and upregulated genes between KO1 and WT from both time points. The dashed line indicates the threshold of q.value = 0.05. **h,** Gene Set Enrichment Analysis (GSEA) enrichment plot of the PI3K-Akt-mTOR signaling pathway. NES, normalized enrichment score; FDR, false discovery rate. **i,** Heatmap of rlog counts of centromeric-kinetochore-associated genes. The asterisk indicates that the gene is differentially expressed. **j,** Comparison of the log_2_ coefficient of variation of nuclear levels of histone marks H4K20me1, H3K9me3 and H3K27me3 in WT and KO1 cells; unpaired two-tailed Wilcoxon test (****, *P*<0.0001). **k,** Comparison of the log_2_ coefficient of variation of nuclear DAPI in WT and KO1 cells; unpaired two-tailed Wilcoxon test (****, *P*<0.0001). Representative images are shown on the right; non-confocal mode, scale bar 25 μm.

**Extended Data Fig. 6:**
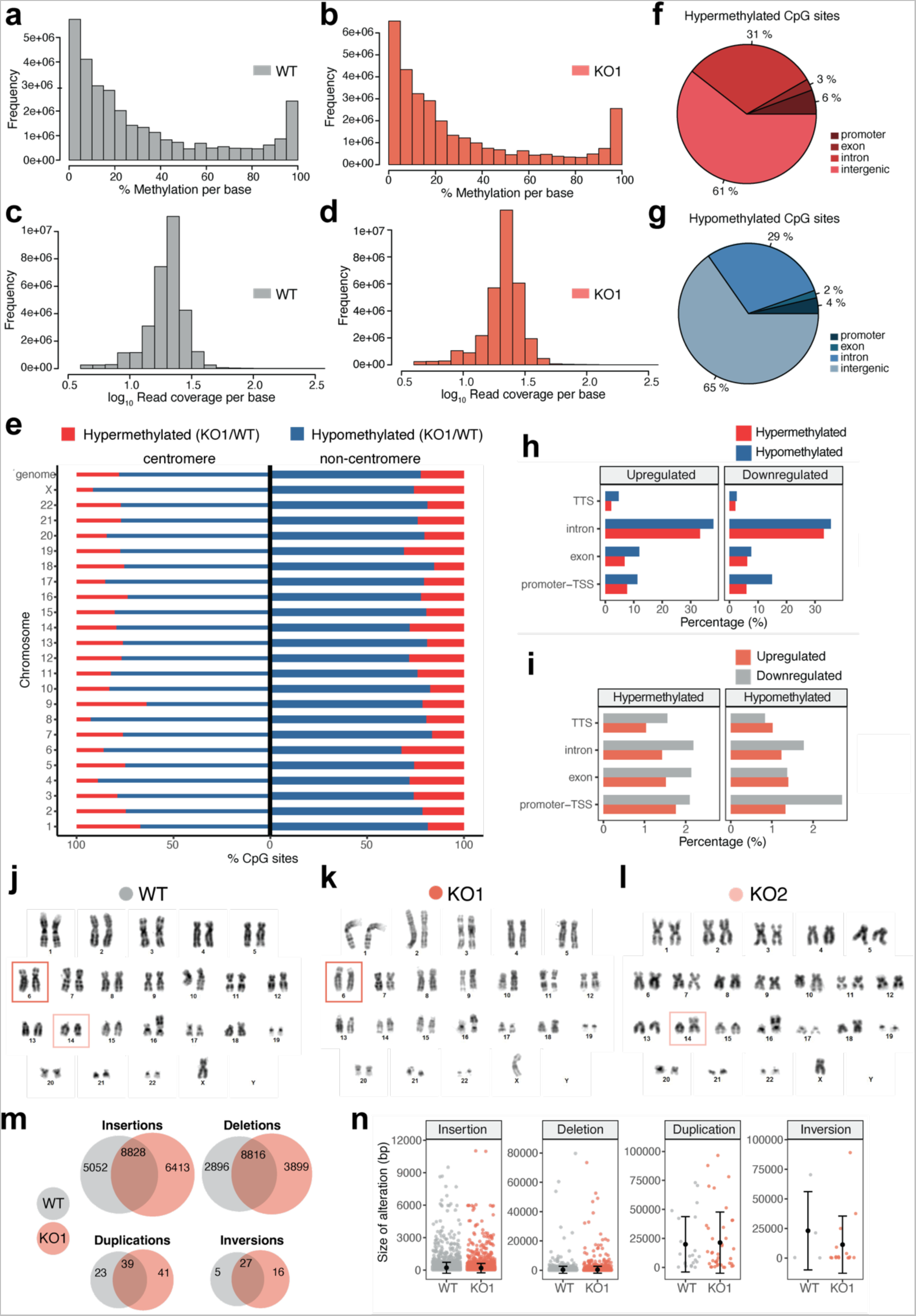
MTHFD2 KO cells show DNA methylation defects and increased structural variation. **a-b**, Histogram showing the frequency of the percentage of methylation per base in HCT116 MTHFD2 wild-type (WT) (a) and knock-out (KO1) (b) cells. **c-d,** Histogram showing the log_10_ read coverage per base in WT (c) and KO1 (d) cells. **e,** Relative percentage of hypermethylated (in red) and hypomethylated (in blue) CpG sites (KO1 versus WT) per chromosome located within centromeric regions (left) or outside centromeric regions (right). **f-g,** Pie charts showing the percentage of hypermethylated (f) and hypomethylated (g) CpG sites found at promoter, exon, intron or intergenic regions. **h,** Percentage of upregulated and downregulated genes with hyper- and hypomethylated CpG sites located in promoter-transcription start sites (TSS), exons, introns or transcription termination sites (TTS). **i,** Percentage of hypermethylated and hypomethylated CpG sites located in up- or downregulated genes, localized in promoter-TSS, exons, introns or TTS. **j-l,** Karyotype of WT (j), KO1 (k) and KO2 (l) cells highlighting the altered chromosomes. As previously described, this male cell line frequently misses the Y chromosome^112^. **m,** Venn diagram of the structural variants detected in WT and KO1 cells. **n,** Size of the structural alterations in base pairs (bp) detected in WT and KO1 cells.

**Extended Data Fig. 7:**
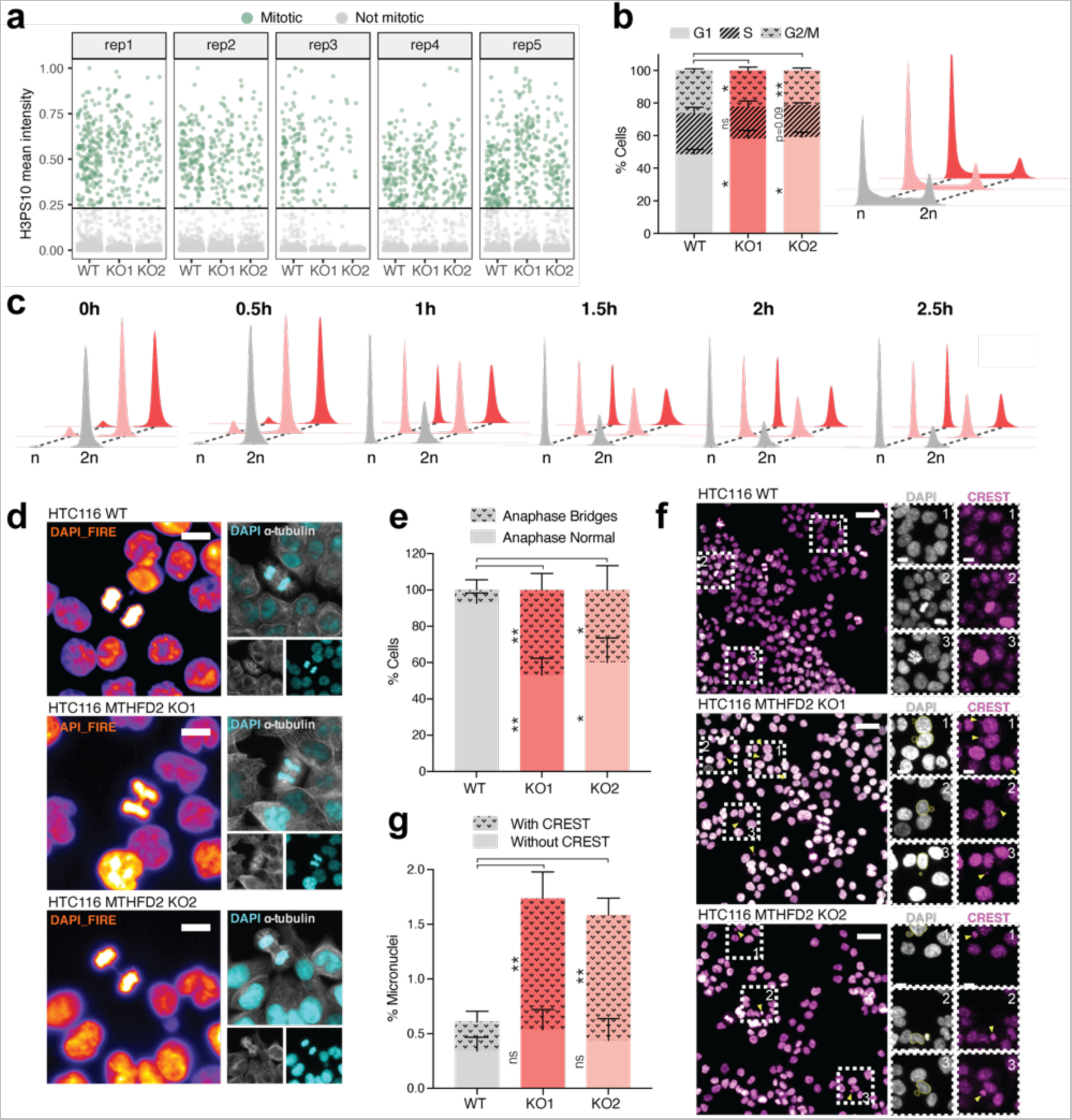
MTHFD2 loss results in reduced mitotic index, mitosis progression delay and mitotic defects. **a**, H3 phospho-Ser10 (H3PS10) scaled mean intensity in HCT116 MTHFD2 wild-type (WT) and knock-out (KO1, KO2) cells. Cells with a signal higher than 3 standard deviations of the average H3PS10 mean intensity were considered mitotic cells. rep, replicate. **b,** Percentage of WT, KO1 and KO2 cells in G1, S and G2/M phases; means + s.d. (*n*=3), unpaired two-tailed *t*-test (left) (ns, non-significant; *, *P*<0.05; **, *P*<0.01). Representative cell cycle profile of WT, KO1 and KO2 asynchronized cells (right). **c,** Representative cell cycle profile of WT, KO1 and KO2 cells at 0, 0.5, 1, 1.5, 2 and 2.5-hour release after RO-3306 drug treatment for 20 hours. **d-e,** Representative images (d) and quantification (e) of anaphase bridges in WT, KO1 and KO2 cells. DAPI is shown in fire (left) or cyan (right) and α-tubulin in grey; scale bar 10 μm. For the quantification, means + s.d. (*n*=3), a minimum of 15 anaphase cells per replicate were analyzed, unpaired two-tailed *t*-test (*, *P*<0.05; **, *P*<0.01). **f-g,** Representative images (f) and quantification (g) of micronuclei in WT, KO1 and KO2 cells. CREST is shown in magenta and DAPI in grey; the scale bar is 20 μm in the global picture and 10 μm in amplified sections. For the quantification, means + s.d. (*n*=3), a minimum of 2400 cells per replicate were analyzed, unpaired two-tailed *t*-test (ns, non-significant; **, *P*<0.01).

**Extended Data Fig. 8:**
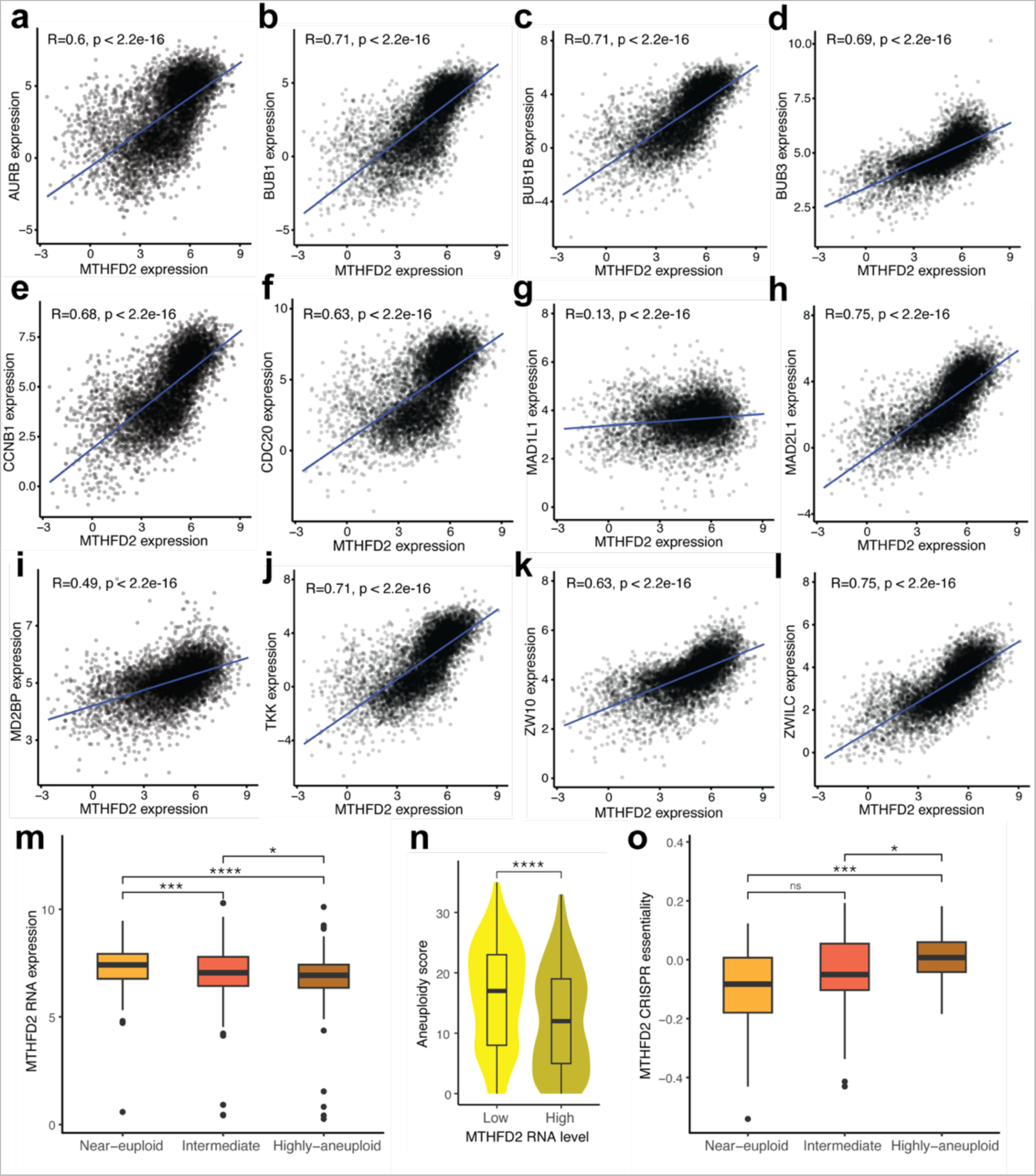
MTHFD2 expression correlates with spindle assembly checkpoint gene expression and aneuploidy score. **a-l**, Correlation between MTHFD2 expression and expression of AURB (a), BUB1 (b), BUB1B (c), BUB3 (d), CCNB1 (e), CDC20 (f), MAD1L1 (g), MAD2L1 (h), MD2BP (i), TKK (j), ZW10 (k) and ZWILC (l). R, coefficient of correlation; P, adjusted p-value. **m,** MTHFD2 RNA expression in the different aneuploidy classification groups of the Cancer Cell Line Encyclopedia (CCLE) cell lines; unpaired two-tailed Wilcoxon test (*, *P*<0.05; ***, *P*<0.001; ****, *P*<0.0001). **n,** Aneuploidy score in CCLE cell lines with the lowest (25% bottom) and highest (25% top) MTHFD2 expression; unpaired two-tailed Wilcoxon test (****, *P*<0.0001). **o,** MTHFD2 CRISPR essentiality in CCLE cell lines with the highest (25% top) MTHFD2 expression in the different aneuploidy classification groups; unpaired two-tailed Wilcoxon test (ns, non-significant; *, *P*<0.05; ***, *P*<0.001).

**Extended Data Fig. 9:**
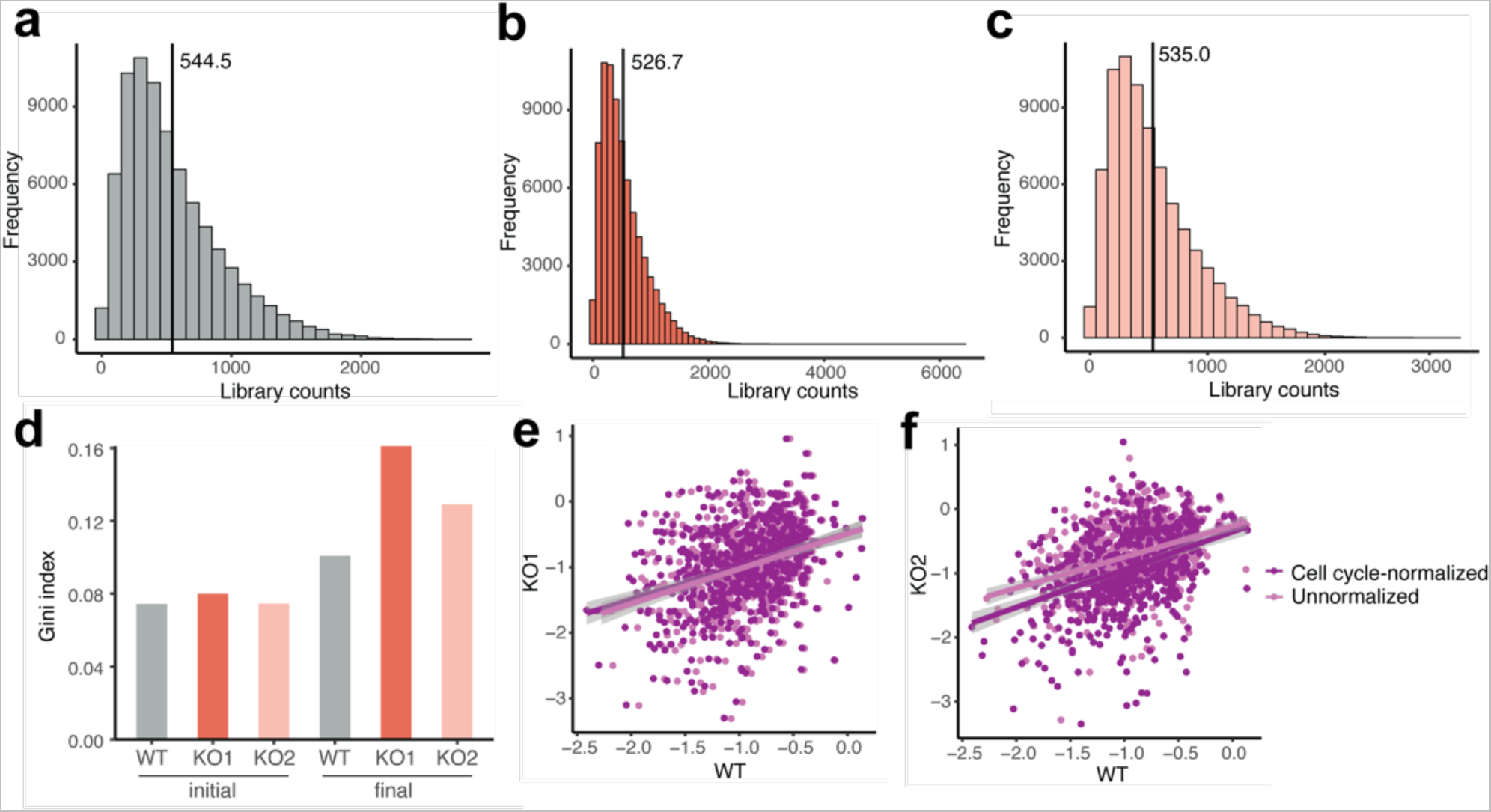
Quality control of genome-wide CRISPR genetic screening. **a-c**, Histograms showing the frequency of library counts in HCT116 MTHFD2 wild-type (WT) (a), knock-out (KO) 1 (b) and KO2 (c) cells. The average library count is indicated. **d,** Gini index of WT, KO1 and KO2 cells at initial and final time points. **e-f,** Correlation of the expression of essential genes between WT and KO1 (e) or KO2 (f) cells before and after cell-cycle normalization.

**Extended Data Fig. 10:**
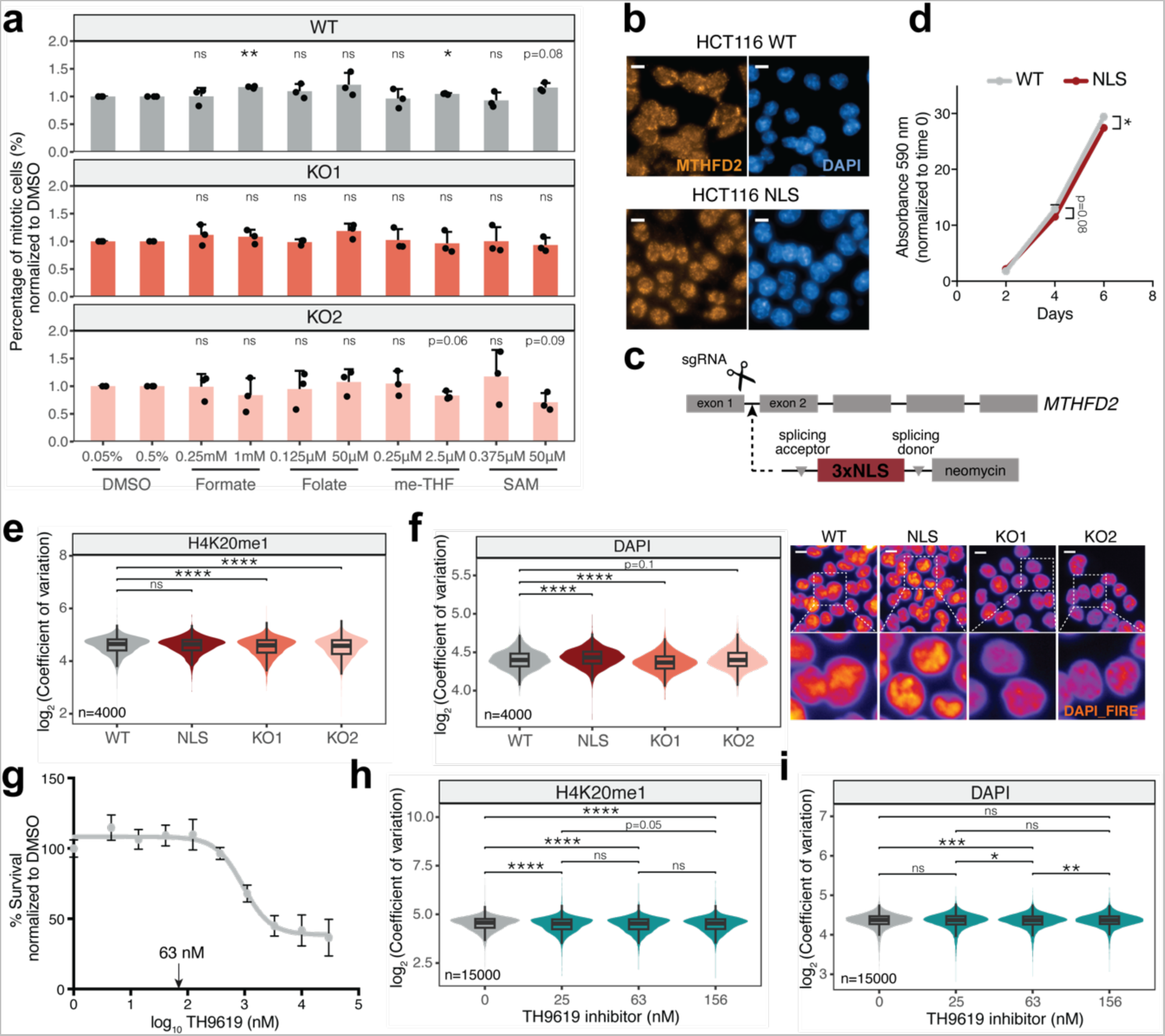
Nuclear MTHFD2 is sufficient for cell proliferation and H4K20me1 methylation. **a**, Percentage of mitotic cells of HCT116 MTHFD2 wild-type (WT) and knock-out (KO1, KO2) after treatment with the indicated concentrations of formate, folate, 5,10- methylenetetrahydrofolate (meTHF), and S-adenosyl methionine (SAM) for 72 hours, and normalized with their respective DMSO condition. The percentage of DMSO corresponds to the highest volume of the low and high metabolite concentrations groups; means ± s.d. (*n*=3), one-sample two-tailed *t*-test (ns, non-significant; *, *P*<0.05; **, *P*<0.01). **b,** Immunofluorescence of HCT116 WT and nuclear (NLS) cells, with MTHFD2 in orange and DAPI in blue; scale bar 10 μm. **c,** Schematic representation of the CRISPR knock-in strategy to introduce the triple nuclear localization signal (NLS) in the MTHFD2 locus. **d,** Growth curve of MTHFD2 WT and NLS cells normalized to time 0 days measured as crystal violet absorbance at 590 nm; means ± s.d. (*n*=3), unpaired two-tailed *t*-test (*, *P*<0.05). **e,** Comparison of the log_2_ coefficient of variation of nuclear levels of H4K20me1 in WT, NLS, KO1 and KO2 cells; unpaired two-tailed Wilcoxon test (ns, non-significant; ****, *P*<0.0001). **f,** Comparison of the log_2_ coefficient of variation of nuclear DAPI in WT, NLS, KO1 and KO2 cells; unpaired two-tailed Wilcoxon test (****, *P*<0.0001). Representative images are shown on the right; non-confocal mode, scale bar 10 μm. **g,** Percentage of survival of HCT116 cells after treatment with the indicated concentrations of the TH9619 inhibitor for 96 hours normalized to the DMSO condition. The 63 nM concentration is indicated. **h-i,** Comparison of the log_2_ coefficient of variation of nuclear levels of H4K20me1 (h) or DAPI (i) in HCT116 cells treated with DMSO or the indicated concentrations of TH9619 inhibitor for 96 hours; unpaired two-tailed Wilcoxon test (ns, non-significant; *, *P*<0.05, **, *P*<0.01; ***, *P*<0.001; ****, *P*<0.0001).

## Supplementary Information

Supplementary Information is available for this paper.

**Supplementary Table 1: All proteins identified in the MTHFD2 pull-down mass spectrometry.**

The first tab contains all the proteins identified in the cytosolic MTHFD2 and IgG immunoprecipitations, and the second tab contains all the proteins identified in the chromatin MTHFD2 and IgG immunoprecipitations. This corresponds to the output of Mascot software.

**Supplementary Table 2: Protein-protein interaction SAINTexpress scores for proteins detected in the MTHFD2 pull-down mass spectrometry.**

The first tab contains the protein-protein interaction scores for all the cytosolic MTHFD2 interactors. The second tab contains the protein-protein interaction scores for all the chromatin MTHFD2 interactors. The third tab contains the list of top nuclear MTHFD2 interactors (fold change >= 5 and Bonferroni False Discovery Rate <= 0.2), along with functional and subcellular localization information.

**Supplementary Table 3: Co-expressed genes of folate metabolism enzymes.**

Each tab contains the list of positively co-expressed genes (r > 0.6 and p-adjusted < 0.05) in at least 10 different cancer types of each enzyme of the folate metabolism.

**Supplementary Table 4: Gene Ontology Biological Process terms enriched among the co-expressed genes of folate metabolism enzymes.**

Each tab contains the list of Gene Ontology Biological Process terms that were significantly enriched (q.value < 0.05) among the list of co-expressed genes of folate metabolism enzymes. This is the output of ClusterProfiler.

**Supplementary Table 5: Differentially expressed genes between MTHFD2 knock-out and wild-type cells.**

The first tab contains the differentially expressed genes (absolute log_2_ fold change > 1.5 and p-adjusted < 0.05) between MTHFD2 knock-out (KO) and wild-type (WT) cells at the initial time point. The second tab contains the differentially expressed genes (absolute log_2_ fold change > 1.5 and p-adjusted < 0.05) between MTHFD2 KO and WT cells at the late time point. The third tab contains the shared differentially expressed genes (absolute log_2_ fold change > 0.58 in at least one time point and p-adjusted < 0.05) between MTHFD2 KO and WT cells in both time points. This corresponds to the output of DESeq2 package.

**Supplementary Table 6: Differentially methylated sites between MTHFD2 knock-out and wild-type cells.**

The first tab contains the list of all differentially methylated CpG sites between MTHFD2 knock-out and wild-type cells. The second tab contains the list of differentially methylated CpG sites located in CpG islands and shores. Those CpG sites with methylation difference >= 15% and q.value < 0.05 were considered differentially methylated. This corresponds to the output of methylKit package.

**Supplementary Table 7: Differentially methylated sites in deleted regions.**

This table contains the information of differentially hypermethylated and hypomethylated CpG sites localized in deletions found in MTHFD2 wild-type and knock-out cells.

**Supplementary Table 8: Synthetic lethal and synthetic viable hits with MTHFD2 knock-out.**

This table contains the list of synthetic lethal hits, with a difference of beta score (KO-WT) < -1, and the list of synthetic viable hits, with a difference of beta score (KO-WT) > 1, along with gene information. This corresponds to the output of MAGeCKFlute.

**Supplementary Methods Table 1: List of plasmids used in the publication.**

**Supplementary Methods Table 2: List of primers and gene blocks used in the publication.**

Correspondence and requests for materials should be addressed to

